# An expanded deep-branching thermophilic bacterial clade sheds light on the early evolution of bacteria

**DOI:** 10.1101/2022.06.14.494929

**Authors:** Hao Leng, Yinzhao Wang, Weishu Zhao, Stefan M. Sievert, Xiang Xiao

## Abstract

The origin of thermophilic bacteria is essential to our understanding of the early evolution of life. However, due to the lack of deep-branching culturable lineages, many controversies remain. Here, we report a novel deep-branching, sulfur-reducing, thermophilic bacterium that was isolated from a deep-sea hydrothermal vent using a newly developed cultivation strategy (“Subtraction-Suboptimal”, StS). This bacterial lineage clusters together with other major thermophilic bacterial groups on phylogenomic trees and forms a robust superphylum-level clade that represents a major, early-diverging bacterial lineage. Ancestral analyses and metabolic modeling reveal the ancestor of this lineage might be a thermophilic and mixotrophic bacteria with a preference for amino acids utilization. These findings provide evolutionary insights into the early divergence of thermophilic bacteria and their adaptive strategies on early Earth.

---

Organisms with optimum growth temperatures higher than 45 °C are designated as thermophiles^1, 2^. These thermophiles are ubiquitously distributed in natural or artificial high-temperature environments, such as terrestrial volcanic sites, hot springs, submarine hydrothermal systems, deep subseafloor, oil reservoirs, solar-heated surface soils and industrial incubators^3–8^. Some thermophiles can also live in alkaline, acidic, or high salinity conditions, highlighting their ability to tolerant multiple stressors^9, 10^. These so-called “extreme environments” are generally considered analogs of conditions existing on early Earth when life originated in the late Hadean or early Archaean eon^11, 12^.

Thermophilic bacteria can be found in a wide range of bacterial phyla, yet only a few phyla contain almost exclusively thermophilic bacteria, namely, Aquificota, Dictyoglomota, Coprothermobacterota, Thermotogota and Thermodesulfobiota. While it has been suggested that thermophily might represent an ancestral feature of bacteria or even life^13–17^, other hypotheses have put forward a non-thermophilic origin of bacteria and placed bacterial lineages such as Chloroflexi or the Candidate Phyla Radiation (CPR) near the root of the bacterial phylogenomic tree^18–20^. A recent study using an outgroup-free gene tree-species tree reconciliation approach rejected previously predicted bacterial roots and placed a new robust root between two major bacterial branches, i.e. Gracilicutes and Terrabacteria, even though with uncertainties regarding the phylogenetic position of the Fusobacteriota lineage^21^. Nevertheless, it still remains unclear when the thermophilic bacteria originated and how they evolved.

The predicted metabolic potential of the last bacterial common ancestor (LBCA) or early diverging bacterial lineage can shed light on the early Earth environments and early ecosystem^17, 22–25^. For thermophilic bacteria, several previous studies focusing on different lineages such as Thermotogota^26^, Aquificota^27^ and Coprothermobacterota^28^ revealed a complex evolutionary history with a high proportion of genes horizontally transferred from other prokaryotes. However, there is still a lack of an evolutionary study based on comprehensive comparative genomic analyses of major thermophilic bacterial clades. Even though cultivation-free metagenomic analyses have greatly expanded our knowledge on bacterial diversity^29, 30^, research on thermophilic bacteria still requires isolated strains to provide direct evidence of their physiology, such as optimal growth temperature to confirm their thermophilic features. Yet, it is extremely difficult to obtain pure cultured novel thermophilic bacteria, especially for deep-branching clades. This strongly limits our understanding of the lifestyle and metabolism of their common ancestor and how these thermophilic bacteria differentiated into new niches.

Here, we report the discovery and isolation of a novel thermophilic, piezophilic, sulfur-reducing bacterial strain, *Zhurongbacter thermophilus* 3DAC. Phylogenomic studies of the strain 3DAC indicate that it closely related to major thermophilic bacterial groups, such as Coprothermobacterota, Dictyoglomota, Caldisericota and Thermotogota. Phylogenetic ancestral analyses indicate that these thermophilic bacterial lineages share one last common ancestor with the potential to metabolize hydrogen, sulfur compounds, and with a mixotrophic carbon catabolism preferring the utilization of amino acids.

## Results and discussion

### Discovery and isolation of a previously unknown thermophilic and piezophilic bacterium

#### Isolation with a novel “Subtraction-Suboptimal” strategy

To increase the probability to cultivate novel thermophilic bacteria from the rare biosphere, we designed a two-stage cultivation strategy, named as “Subtraction-Suboptimal” (StS) strategy (Fig. 1c). The concept of this method is to reduce the complexity of the community and minimize the abundance of dominant heterotrophic organisms by using first an autotrophic culture medium and subculturing for several times (denoted “Subtraction”), followed by switching to a rich heterotrophic medium at a suboptimal temperature to recover the target heterotrophic thermophilic bacteria present in low abundance and to avoid competition from usually dominant fast-growing hyperthermophilic heterotrophic archaea (denoted “Suboptimal”).

**Fig. 1.**
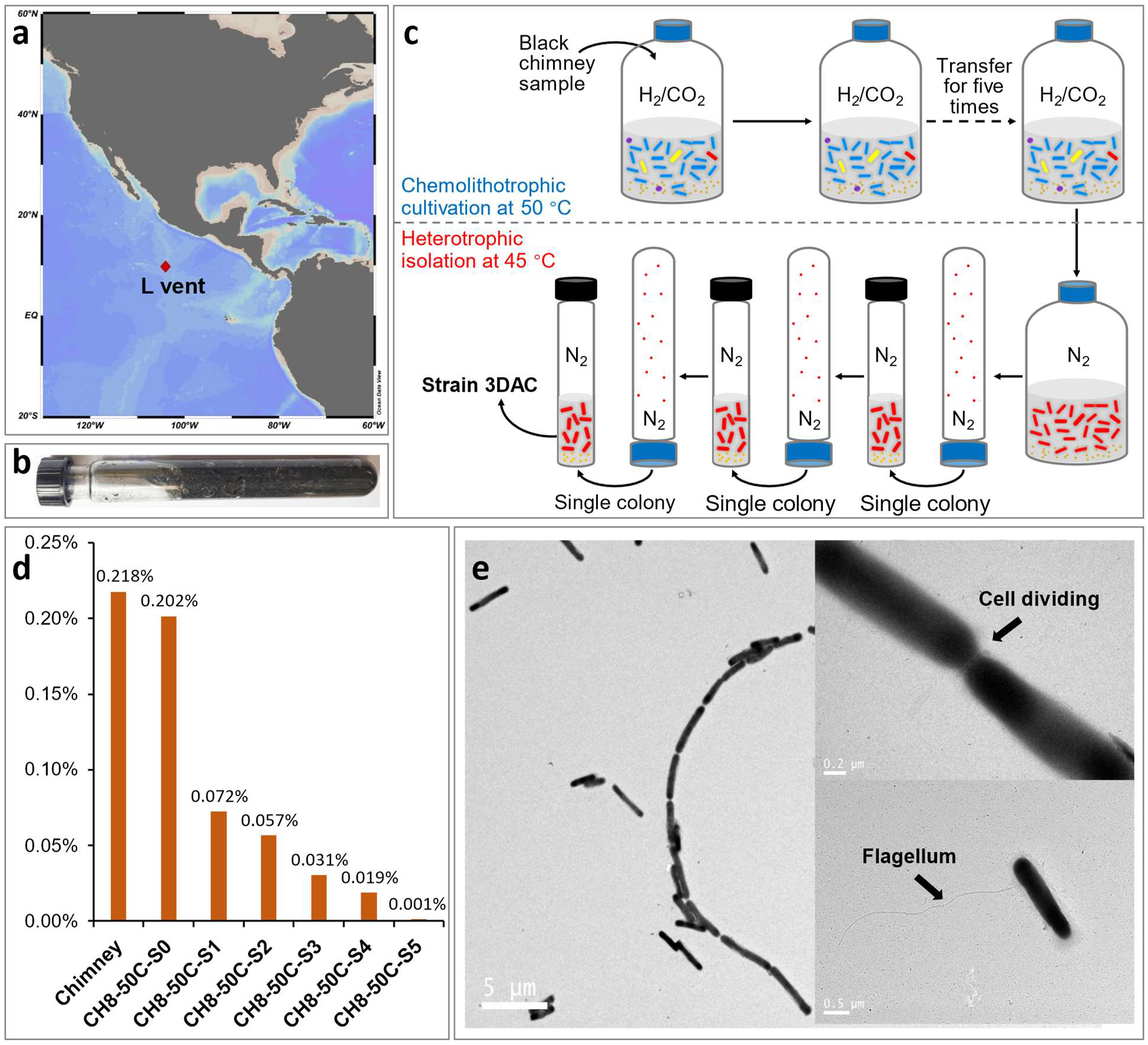
Strain isolation procedure. **a**, Location of L vent at the 9°N vent field on the East Pacific Rise. **b**, The black L vent chimney sample collected in an anaerobic tube. **c**, Schematic diagram of the “Subtraction-Suboptimal” (StS) strategy used to isolate *Zhurongbacter thermophilus* strain 3DAC in this study. **d**, The track of proportion changes of 3DAC closely related sequences in the chimney sample, original enrichment, and different transfers. Labels: Chimney, the chimney sample; CH8-50C-S0, original enrichment; CH8-50C-S1 to -S5, the first to fifth subcultures transferred from the original enrichment. All samples indicate that strain 3DAC represents a rare species. **e**, Transmission electron micrograph of 3DAC, in which 3DAC cells form long filaments with chains of up to 11 cells or more; the micrograph showing a connection between two 3DAC cells; and a long lateral flagellum of 3DAC.

In this case, the original sample, collected from an active black smoker chimney emitting hydrothermal fluid with a temperature of 231 °C (Figs. 1a, b; Hou et al., 2020^31^), was firstly cultivated in a chemolithoautotrophic medium (SME medium, see Methods) at 50 °C, then consecutively transferred for five times under autotrophic condition, reducing the microbial community complexity and the ratio of heterotrophs. The conditions of the chemoautotrophic “Subtraction” stage mimicked the natural environment *in situ* and established a community dominated by autotrophs (mainly *Hydrogenimonas* within Campylobacterota and *Desulfurobacterium* within Aquificota)^32^. It performed similarly to the traditional enrichment method by decreasing the proportion of heterotrophs, but kept the co-cultured heterotrophs with autotrophs. In general, while the community is simplified through the “Subtraction” stage, it still maintains a certain degree of diversity, providing more opportunities for the isolation of rare species supported by autotrophs.

For the “Suboptimal” stage, we switched the cultivation condition from the autotrophic medium to a heterotrophic medium to provide a strong selection for heterotrophs. However, we decreased the cultivation temperature from 50 °C to 45 °C. While this resulted in a suboptimal growth temperature for thermophilic bacteria, it prevented the growth of heterotrophic hyperthermophilic archaea (i.e., *Thermococcus,* growth temperature ranges between 50 and 100 °C^33^), which were pervasive in the chimney sample, allowing the recovery of rare heterotrophic thermophilic bacteria. After a four-day incubation, growth was observed, followed by the isolation of single transparent and round colonies by using the rolling tube method^34^ containing solid TRM medium.

During the “Subtraction” stage, some low abundance heterotrophic species co-cultured with the autotrophs can be retained, these rare species can be hardly isolated by traditional methods when fast-growing heterotrophic microorganisms present at the same time. The successful isolation of the strain 3DAC, which represented only ∼0.001% of the total community in the fifth transfer of the chemoautotrophic enrichment (∼0.2% in chimney sample, Fig. 1d), highlights the advantage of the “StS” strategy to isolate rare species from environmental samples.

#### Morphology and physiology

The cells of strain 3DAC are rod-shaped, 1.3-7.5 μm long (average 3.1 ± 1.3, *n* = 25) and 0.4-0.6 μm wide (average 0.5 ± 0.07, *n* = 25). They have a single lateral flagellum and are highly motile under a light microscope. During growth, the strain 3DAC occurs mostly as single cells, but sometimes forms long chains (Fig. 1e, Extended Data Fig. 1). The strain 3DAC grows only under strictly anaerobic conditions and performs as a piezophilic thermophile: temperature range of 30–75 °C (optimal 70°C), NaCl range of 1.0-4.5% (w/v) (optimal 2.5 %), pH range of 5.5-8.5 (optimal pH 7.0), and hydrostatic pressure range of 0.1-80 MPa (optimal 20 MPa) (Extended Data Figs. 2a-d).

**Fig. 2.**
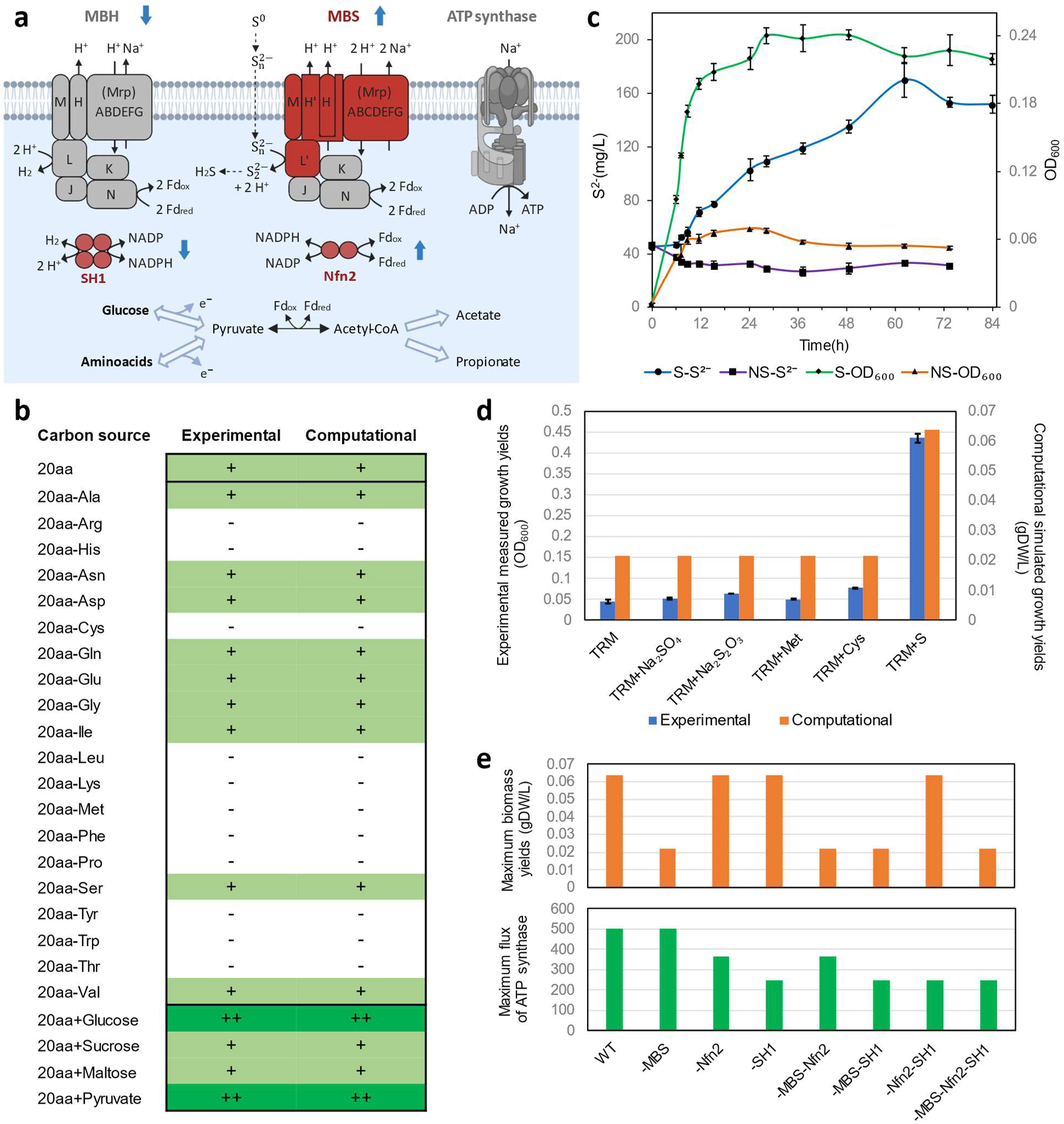
Energy metabolism of strain 3DAC. **a**, Schematic diagram of the energetic process in strain 3DAC. Red shapes represent archaeal orthologs. Gray shapes represent bacterial orthologs. Solid lines represent enzymatic reactions. Dotted lines represent abiotic reactions. Blue arrows indicate significant up- or down-regulation with sulfur in the culture medium based on transcriptomic data [log2(fold change) > 1, p-value < 0.05]. **b**, Growth capabilities on varied carbon sources of strain 3DAC. Computational predictions were simulated by the genome-scale metabolic model of 3DAC (Supplementary Data 1). “20aa”: all 20 amino acids supplemented as sole carbon source. “20aa-X”: removal of individual amino acids X. “20aa+Y”: extra carbon source Y supplied with 20 amino acids. “+”: similar growth yields compared with the 20aa condition; “++”: at least 1.5-fold higher growth yields than 20aa; “-”: no growth. **c**, Sulfide concentration and OD_600_ during the growth of strain 3DAC. S: medium with elemental sulfur; NS: no elemental sulfur. **d**, Experimental and computational growth yields of the 3DAC strain by utilizing various sulfur species. (**c** and **d**) The error bars represent standard deviations. **e**, Simulations of wild-type 3DAC (WT), single deletions (-MBS, -Nfn2, -SH1), double deletions (- MBS-Nfn2, -MBS-SH1 and -Nfn2-SH1) and triple deletions (-MBS-Nfn2-SH1) of archaea-derived complexes in energetic processes at 3DAC.

#### Carbon source utilization

Separately adding peptone, casamino acids, beef extract, yeast extract, and tryptone at a final concentration of 0.2% (w/v) supported the rapid growth of strain 3DAC (Extended Data Figs. 2e). In a basal TRM medium without a carbon source, no growth was detected upon separately adding only one type of any carbohydrate or organic acid (see Methods) under the tested conditions. However, strain 3DAC grew well with 20 canonical amino acids simultaneously added to the basal TRM medium. Moreover, “leave-one-out” experiments with 20 amino acids revealed that 3DAC relied on 11 essential amino acids (i.e. L-arginine, L-cysteine, L-histidine, L-leucine, L-lysine, L-methionine, L-phenylalanine, L-proline, L-threonine, L-tryptophan and L-tyrosine) (Fig. 2b), no growth was observed as the absence of any one of these amino acids. In the presence of the 20 amino acids, D-(+)-glucose and pyruvate could be used as additional carbon sources (Fig. 2b, Supplementary Data 1).

### Genomic and phylogenetic analyses indicate 3DAC as a deep-branching, sulfur-reducing heterotrophic bacterium

#### Genome characteristics

The complete genome of strain 3DAC is a single circular chromosome of 1,588,679 bp with a GC content of 41.15% (Supplementary Fig. 1). The 3DAC genome contains three rRNA gene operons and 46 tRNA genes. A total of 1,524 coding DNA sequences (CDSs) with an average length of 980.78 bp, covering 94.09% of the overall genome, can be identified in the genome. The strain 3DAC has a complete glycolysis pathway, suggesting a capability of carbohydrate utilization, but many genes related to amino acid synthesis are absent, indicating the requirement of certain amino acids. This phenomenon is consistent with the results of physiological experiments, the growth of 3DAC depends on the addition of amino acids and can be stimulated by adding carbohydrates (Fig. 2b). The loss of genes related to amino acid synthesis is common among thermophiles. Since the synthesis of amino acids consumes much energy^35^, it is more efficient for thermophiles to absorb amino acids directly from the environment, which is more conducive to their adaptation to the hydrothermal environment.

#### Archaea-derived energetic process in 3DAC

The respiratory energetic process in strain 3DAC contains a Na^+^-driven ATP synthase (Extended Data Fig. 3), a 12-gene cluster coding a membrane bound hydrogen-producing complex (MBH) and a 13-gene cluster coding the membrane bound sulfur respiration complex (MBS), which is similar to the energetic process in hyperthermophilic archaea within the order Thermococcales^36–38^ (Fig. 2a). MBS enables cells to reduce elemental sulfur (S^0^) to hydrogen sulfide (H2S) *in vivo* and to use reduced ferredoxin as an electron donor^37^. Genes in MBH of 3DAC, including the key catalytic subunit of hydrogen production (MbhL), are bacterial orthologs, as expected, while most of the MBS-coding genes (10 of 13) of 3DAC are archaeal orthologs encoding the complete membrane arm (MbsA, B, C, D, E, G, H, H’ and M subunits) and the key catalytic subunit of sulfur reduction (MbsL) (Fig. 2a, Supplementary Data 1). Compared to the bacterial hydrogen-producing MBH, the archaea-derived MBS enable cells to convert energy more efficiently with a 3.4-fold higher redox potential to reduce elemental sulfur (S^0^) to hydrogen sulfide (H2S) than MBH^36, 37^. In addition, two redox-balancing complexes in 3DAC, soluble hydrogenase I (SH1) and NADP-dependent ferredoxin oxidoreductase II (Nfn2), are also archaeal orthologs with a high identity (>60%) as genes reported in hyperthermophilic archaea Thermococcales^39, 40^.

#### Sulfur utilization

Consistent with genomic analysis, physiological experiments revealed that the presence of elemental sulfur significantly stimulated the growth of strain 3DAC, compared with the medium without sulfur (Fig. 2c, d). Other sulfur substitutes, such as sodium sulfate, sodium thiosulfate, L-cysteine or L-methionine, did not significantly stimulate growth (Fig. 2d, Supplementary Data 1). During the growth of 3DAC, elemental sulfur is reduced to sulfide (Fig. 2c). This process is proposed to be conducted by MBS with the help of redox balancing via Nfn2 and/or SH1, which is also in agreement with transcriptomic analysis with or without elemental sulfur. We found that most genes (10 of 13) in MBS were significantly upregulated with elemental sulfur in the medium, including the catalytic subunit MbsL. Genes in Nfn2 were also highly expressed with sulfur, indicating the potential coupling of Nfn2 with MBS for redox balancing. In contrast, genes in MBH and SH1 were significantly downregulated in the presence of sulfur, which suggested competition between MBS and MBH and potential coupling between MBH and SH1 (Supplementary Data 1).

#### Necessity of archaea-derived MBS, SH1 and Nfn2 for growth and ATP production

A genome-scale metabolic model of 3DAC, GEM-i3DAC (Supplementary Data 1), was constructed to simulate the impacts of the respiratory and redox balancing complexes derived from archaea (i.e., MBS, SH1 and Nfn2) in 3DAC. The model incorporated both annotations from automatic pipelines and manually curated biochemical evidence of (hyper)thermophiles from the literature (Supplementary Data 1). This model was validated to predict growth trends accurately by utilizing varied carbon sources and electron acceptors compared to the experimental observations (Figs. 2b, d). Simulations were further performed on the 3DAC wild-type, single, double, and triple deletions of the archaea-derived complexes MBS, SH1 and Nfn2. Deletion of MBS blocked sulfur consumption and hydrogen sulfide production and theoretically decreased maximum biomass yields to 33% of the WT (Fig. 2e, Supplementary Data 1), which supported the necessity of MBS for utilizing sulfur as an electron acceptor to stimulate 3DAC growth. The single deletion of Nfn2 and SH1 decreased the maximum flux of ATP synthase to 72% and 50%, respectively, which indicated the essential function of Nfn2 and SH1 in ATP production. These results revealed the necessity of these archaea-derived complexes for both growth and ATP production in the 3DAC strain. Overall, strain 3DAC performed as a thermophilic heterotroph combining bacteria-derived hydrogen-producing and archaea-derived sulfur-reducing complexes for respiration.

#### Phylogenetic analyses identify 3DAC as a deep-branching lineage

Phylogenomic analysis based on different concatenated conserved protein sequences^21, 41–43^ and 16S rRNA gene-based analysis both place the phylogenetic position of strain 3DAC as a sister lineage to Coprothermobacterota, a bacterial phylum recently proposed by Pavan *et al*. (2018)^44^ (Figs. 3a-c, Extended Data Fig. 4a-c). However, 3DAC falls separately from all Coprothermobacterota strains, and the average amino acid identity (AAI) shared by these strains was only 48-49% (Supplementary Table 1). The most closely related cultured strain of 3DAC based on both 16S rRNA gene phylogenetic and phylogenomic analyses was *Coprothermobacter proteolyticus* DSM5265. Nevertheless, the 16S rRNA gene sequence (three copies) identities between 3DAC and DSM5265 are only 81.25-83.00% (Supplementary Table 2), and they also have large physiology differences. *C. proteolyticus* DSM5265 is an anaerobic thermophilic microbe frequently identified in artificial thermophilic anaerobic systems, it grows on various proteinaceous substrates and carbohydrates. The strain DSM5265 grow better without sodium chloride and are able to reduce thiosulfate to sulfide^28, 45–47^, while 3DAC was isolated from natural hydrothermal vent and has a preference to utilize amino acid and elemental sulfur. Phylogenetic analysis of the 16S rRNA genes shows that strain 3DAC, together with other related 16S rRNA genes from enrichment cultures from tubes of *Alvinella pompejana* collected from a deep-sea vent at 13°N on the East Pacific Rise, forms a monophyletic cluster, clearly separating them from Coprothermobacterota (Extended Data Fig. 3c). We also compiled almost complete 16S rRNA gene sequences (longer than 1400 bp, > 95% coverage) from Coprothermobacterota and compared the sequence identities with those of the 3DAC clade. The 16S rRNA gene sequence identities ranged from 75.73% to 83.00% (Supplementary Table 3). Since strain 3DAC was isolated from a high-temperature hydrothermal vent environment, we propose 3DAC as *Zhurongbacter thermophilus* 3DAC, named after Zhurong, the fire god in Chinese myth. For higher taxonomic classification, based on the suggested AAI ranges for the whole-genome^48^ and median 16S rRNA gene sequence^49^ identities to distinguish a new phylum, it is highly possible that this lineage represents a previously unknown bacterial phylum. However, the RED (relative evolutionary divergence) score of strain 3DAC is 0.5456, indicating that 3DAC might belong to a class-level lineage within the phylum Coprothermobacterota based on GTDB taxonomy^50^. There is a discordance between traditional classification and GTDB, hence we only temporally call the clade represented by 3DAC as a deep-branching lineage Zhurongbacterota until more 3DAC related species are discovered for more accurate taxonomic assessment, as well as the prokaryotic taxonomic definition of phylum reach a consensus within scientific community.

**Fig. 3.**
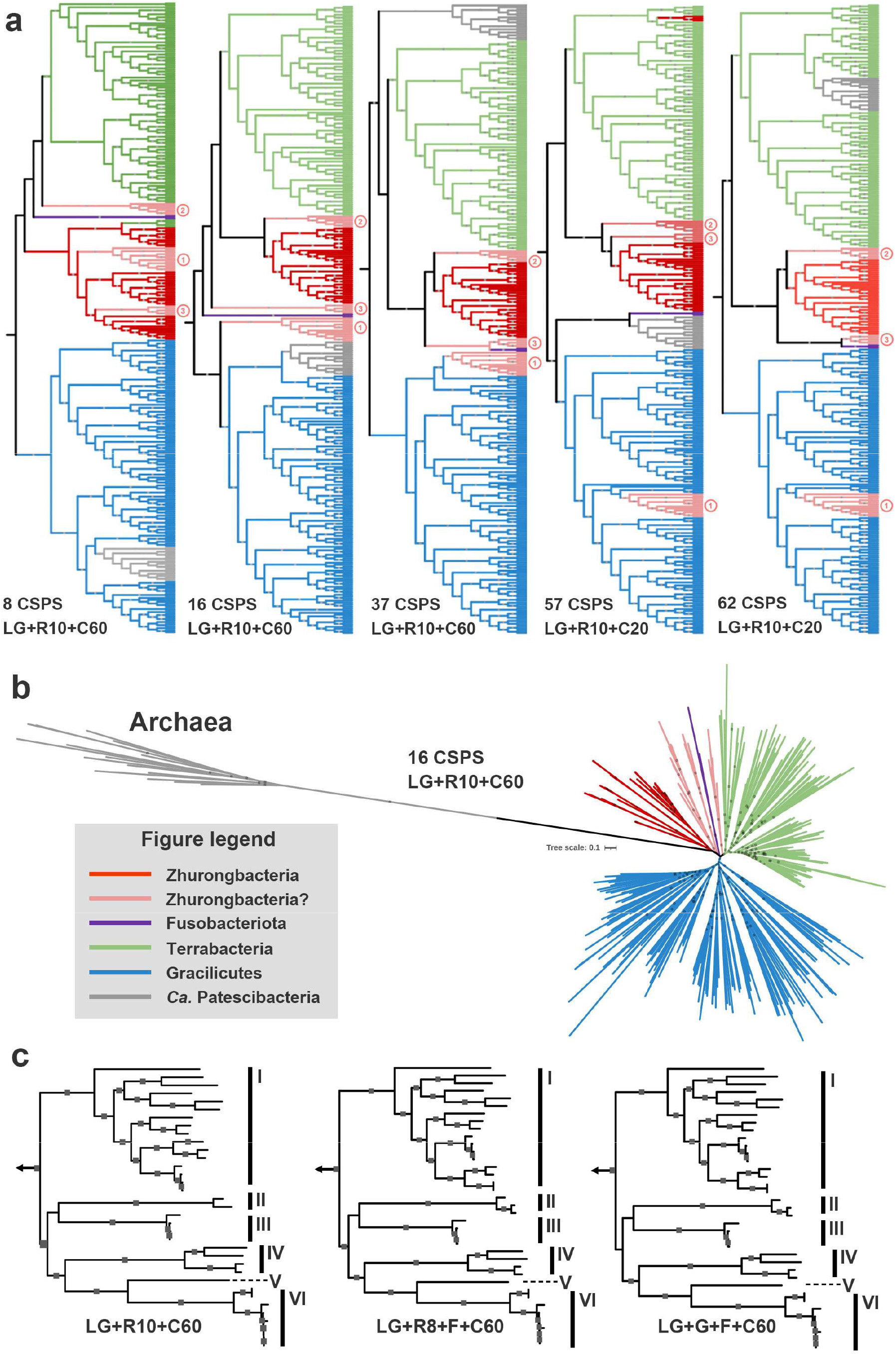
Phylogenetic analyses. **a**, Phylogenomic tree of *Zhurongbacter thermophilus* 3DAC and other major thermophilic bacteria using 8/16/37/57/62 conserved protein sequences. Red color represents major thermophilic bacterial phyla, namely, Zhurongbacter, Caldisericota, Coprothermobacterota, Dictyoglomota, Thermotogota, and Thermodesulfobiota. The pink color with circled numbers refers to Aquificota (one), Synergistota (two) and Deinococcota (three), while the purple color refers to Fusobacteriota. The CPR lineage here is in grey color. **b**, Phylogenomic tree of *Zhurongbacter thermophilus* 3DAC and other major thermophilic bacteria using Archaea as the outgroup with 16 conserved single-copy protein sequences (CSCP) with IQ-Tree using the LG+R10+C60 model. **c**, Phylogenomic inferences of the lineages from the superphylum Zhurongbacteria with 37 conserved protein sequences with different models but without Archaea genome. I-VI represent Thermotogota, Thermodesulfobiota, Dictyoglomota, Caldisericota, Zhurongbacter and Coprothermobacterota. The arrows indicate other bacterial branches.

### Evolutionary adaptation of the thermophilic superphylum Zhurongbacteria

#### Major deep-branching thermophilic bacterial phyla cluster closely together on the phylogenomic tree

Phylogenetic analysis based on different sets of conserved protein sequences and using different inference models revealed that, in addition to Coprothermobacterota^28, 44, 47^, 3DAC is most closely related to other major thermophilic bacterial groups, i.e. Caldisericota^51, 52^, Dictyoglomota^53^, Thermotogota^13, 54, 55^, Bipolaricaulota^56, 57^ and Thermodesulfobiota^41, 44, 58–61^ (Fig. 3a-c). These thermophilic bacterial lineages robustly clustered together, forming a monophyletic lineage supported by phylogenomic analyses as previously reported^41^ (Fig. 3a-c). We also tested their phylogenetic positions with two outgroup rooting datasets, i.e. using Archaea as an outgroup but without CPR branch that would influence the phylogenetic inference when considering archaeal lineages^21^, and using bacterial root position proposed by Coleman et al. (2021)^21^. The results indicate that these thermophilic groups consistently cluster near the root of Archaea-Bacteria phylogenomic tree, or display a deep-branching monophyly with the sister clade Terrabacteria by using the two rooting methods, respectively (Fig. 3a, b), representing a major, early-diverging bacterial lineage. Moreover, all the above-mentioned bacterial lineages also share similar metabolic capabilities; for example, having an anaerobic lifestyle; nearly all their lineages are able to use elemental sulfur or sulfur-related compounds; most members are heterotrophs with the ability to degrade sugars or/and proteins (Supplementary Table 4). Therefore, here we propose a superphylum-level clade, Zhurongbacteria, for thermophilic bacterial phyla that cluster closely with 3DAC and branch together near the bacterial root on phylogenomic trees. This thermophilic superphylum-level clade contains at least six lineages that have pure cultured strains: Zhurongbacter, Caldisericota, Coprothermobacterota, Dictyoglomota, Thermotogota, and Thermodesulfobiota. There are also several other candidate phyla cluster with the Zhurongbacteria clade near the root of bacterial 16S rRNA gene tree but display some inconsistence on the phylogenomic tree using different conserved marker protein sequence sets, such as thermophilic Aquificota. We have carefully checked the phylogeny of Aquificota by different methods, and found that most of genes within Aquificota have undergone horizontal gene transfer (HGT) as previously described^62, 63^ (Extended Data Fig. 5, 6). With increasing number of conserved marker protein sequences for constructing the phylogenomic trees (from 8 to 62 marks), the phylogenetic position of Aquificota moved from Zhurongbacteria away to Campylobacterota or Proteobacterota (Figure 3a), suggesting a potential influence by HGTs.

Some other candidate phyla such as Deinococcota and Synergistota also cluster with the Zhurongbacteria clade (Figure 3a) and therefore might belong to Zhurongbacteria. However, although many of their members are also thermophilic bacteria, yet these two phyla also contain various non-thermophilic bacteria^64–66^. Further study regarding whether the clades Deinococcota and Synergistota had a thermophilic origin need to be resolved.

#### Last common ancestor of Zhurongbacteria and its evolutionary adaptation

Phylogenetic ancestral analyses indicate that the members of the thermophilic clade Zhurongbacteria (Zhurongbacter, Caldisericota, Coprothermobacterota, Dictyoglomota, Thermotogota, and Thermodesulfobiota) might have shared one potentially last thermophilic bacterial common ancestor (LTBCA). The predicted ancestral gene set from six Zhurongbacteria lineages, in general, includes genes coding for thermophilic-related components, a near complete flagellar assembly that enables its movement, hydrogenase and putative sulfur metabolism for energy conservation, as well as the Wood-Ljungdahl pathway and saccharide and amino acid utilization pathways (Supplementary Data 2). Nevertheless, the amino acid biosynthesis pathways are not complete, possibly due to the limitation of ancestral prediction (Fig. 4a). A genome-scale metabolic model of LTBCA was further constructed and suggests it is a motile mixotrophic bacterium with carbon dioxide, saccharides and proteins as substrates (Supplementary Data 2). This predicted ancestral metabolism of the superphylum Zhurongbacteria also shares many similarities with the recently inferred metabolic capacities of last universal common ancestor (LUCA) and LBCA^17, 21, 67^. LTBCA, LUCA and LBCA all possess a Wood-Ljungdahl pathway and are considered strictly anaerobic microorganisms. LTBCA and LUCA are predicted both thermophilic and share the potential for metabolizing hydrogen, while LBCA and LTBCA share the predictions for motility and the potential for degrading organic carbon compounds. From these ancestral reconstructions, the prediction of their pathways clearly shows a gradual increase in their metabolism inventories during the potential evolutionary transgression from LUCA to LBCA and then LTBCA (Extended Data Fig. 7).

**Fig. 4.**
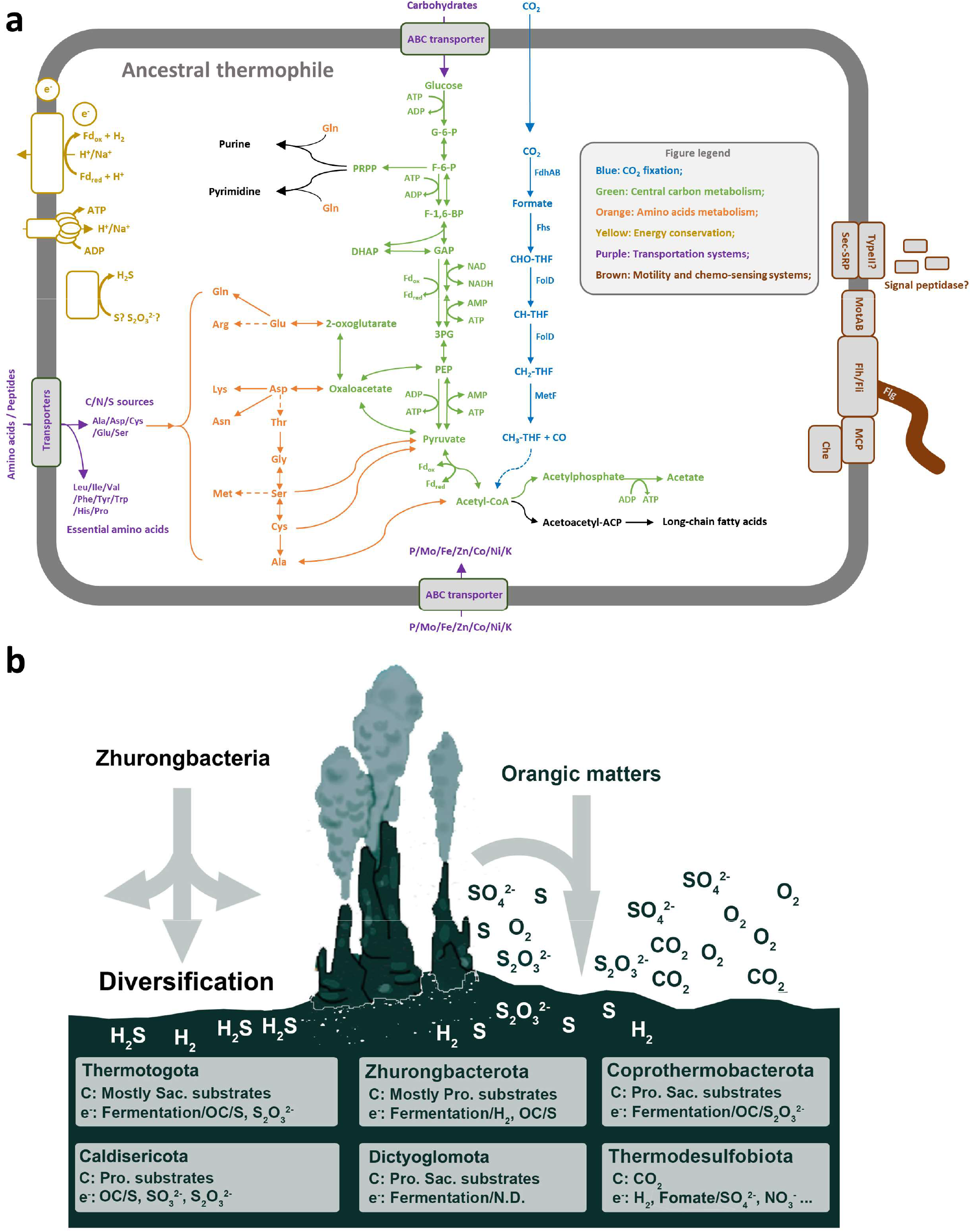
Metabolic capability of the Zhurongbacteria ancestor and its diversification. **a,** Ancestral metabolic diagram of the thermophilic superphylum Zhurongbacteria. The ancestral gene set prediction indicates that the thermophilic bacteria ancestor may prefer a movable heterotrophic lifestyle with hydrogenase but also contains a Wood-Ljungdahl pathway. b. Ecological niches of the thermophilic superphylum Zhurongbacteria. Illustration of metabolic feathers of different bacterial lineages from the thermophilic superphylum Zhurongbacteria (Supplementary Table 4), the hydrothermal vent here indicates that these bacteria might have originated from similar environments on early Earth but under adaptation radiation during their evolutionary history. Pro. refers to proteins, Sac. refers to saccharides, OC refers to organic carbon.

After the diversification of the thermophilic Zhurongbacteria ancestor, each lineage adapted to different environments and gradually evolved to form specific features by gene gain and loss (Fig. 4b, Extended Data Fig. 8), especially on sulfur metabolism and carbon utilization pathways. For sulfur respiration, most of the Zhurongbacteria contained the putative catalytic subunit MbsL-encoding gene of the MBS complex-encoding gene cluster (Extended Data Fig. 9)^37^, while Coprothermobacterota, Zhurongbacter and Thermotogota also contain the catalytic subunit MbhL-encoding gene of the MBH complex-encoding gene cluster, coding for a homologous enzyme complex responsible for hydrogen metabolism rather than sulfur. With the combination of both MBS and MBH complex within cells, microorganisms could gain more environmental fitness, as revealed by the sulfur respiration experiments and transcriptome of *Zhurongbacter thermophilus* 3DAC. As the sulfur respiration capability strongly stimulates the growth in 3DAC here and other strains in previous studies^68^, it indicates that their ancestor might originate from a sulfur compound-rich hydrothermal vent or hot spring environment. Indeed, nearly all of the descendants of the Zhurongbacteria ancestor are still active in diverse hydrothermal ecosystems worldwide^69, 70^. For carbon assimilation, even though the predicted Zhurongbacteria ancestor was potentially able to use carbon dioxide, saccharides and proteins as substrates, members of the clades Zhurongbacter, Caldisericota, Coprothermobacterota, Dictyoglomota and Thermotogota are considered thermophilic heterotrophic bacteria and can use saccharides, proteins or both as substrates, whereas members of the Thermodesulfobiota are considered autotrophic bacteria and fix carbon dioxide during growth. Nevertheless, the metabolic reconstruction shows that the ancestor of the Zhurongbacteria was a mixotrophic bacterium that was capable of both heterotrophic and autotrophic growth under different conditions. Some of the descendants of this Zhurongbacteria ancestor, such as the phyla Coprothermobacterota and Thermotogota, may have lost their autotrophic ability during adaptation to a heterotrophic lifestyle, while Thermodesulfobiota gained an additional autotrophic strategy. Those thermophilic heterotrophic bacteria in the superphylum Zhurongbacteria also diversified their preferences for carbon assimilation (Extended Data Fig. 8). For example, the evolutionary histories of members in the phyla Dictyoglomota and Caldisericota are similar. They both gained a number of genes for peptide and sugar transport and several energy conservation genes, while lost their motility genes. The lineage Zhurongbacter discovered here also gained peptide and sugar transport genes. The evolutionary histories of the phyla Coprothermobacterota and Thermotogota are similar, as they both gained genes for sugar transport, but lost some of the genes for biosynthesis of amino acids, and both of these groups are frequently discovered in artificial high-temperature facilities such as industrial reactors while many bacterial strains from Thermotogota have already been widely used in synthetic biology^13, 28, 47^.

In conclusion, there is a clear evolutionary path by which these bacteria adapted to different niches with preferences for carbon dioxide, proteins or saccharides as carbon substrates (Fig. 4b). As cultivation and sequencing technologies are developing rapidly, more deep-branching thermophilic bacteria from the superphylum Zhurongbacteria will be confirmed or discovered, and with more advanced phylogenetic inference methods, the evolutionary history of this mysterious bacterial lineage may become much clearer, even shedding further light on the metabolic properties of the last universal common ancestor.

## Materials and Methods

### Sample collection

The original sample was collected with the remotely operated deep-sea research vehicle (ROV) *Jason II* during R/V *Atlantis* research cruise AT26-10 (Dive 761) from a black smoker chimney at L-vent located at the 9°N deep-sea vent field on the East Pacific Rise (9°46.25’ N, 104°16.74’ W, water depth 2,528 m). The temperature of the emitted fluid was 231 °C. More information about the fluid geochemistry and microbial communities in the chimney walls can be found in Hou *et al*. (2020)^31^. Pieces of the chimney were stored immediately upon retrieval in sterile anaerobic tubes, and the headspace in the tubes was filled with pure nitrogen gas with slight overpressure to prevent the entry of oxygen. The samples were then stored at 4 °C until enrichment in the laboratory.

### “Subtraction-Suboptimal” strategy guided isolation

The outer layer of the black chimney sample (0.25 g) was first enriched anaerobically in a SME medium^71^, which contained the following components (L^−1^): NaCl, 20.0 g; MgSO_4_·7 H_2_O, 3.5 g; MgCl_2_·6 H_2_O, 2.75 g; KCl, 0.325 g; KNO_3_, 2.0 g; MES, 4.27 g; NaBr, 50.0 mg; H_3_BO_3_, 15.0 mg; SrCl_2_·6 H_2_O, 7.5 mg; (NH_4_)_2_SO_4_, 10·0 mg; KI, 0.05 mg; Na_2_WO_2_·2H_2_O, 0.1 mg; CaCl_2_·2H_2_O, 0.75 g; KH_2_PO_4_, 0.5 g; NiCl2·6H_2_O, 2.0 mg; and resazurin, 1.0 mg. The pH of the medium was adjusted to 6.0 using 1 M NaOH. After autoclaving, 1 ml of trace element mixture, 1 ml of vitamin mixture, 1 ml of thiamine solution, and 1 ml of vitamin B12 solution^72^ were added to the medium (L^−1^). Then, 50 ml aliquots of the medium were separated into 150-ml glass serum bottles and supplemented with 0.5 g of elemental sulfur and 0.5 ml of Na_2_S·9H_2_O (10 %, w/v; pH 7.0), and the tightly stoppered bottles were pressurized with hydrogen/carbon dioxide (80: 20; 101 kPa). The cultures were incubated at 50 °C. We obtained the original enrichment in this chemolithotrophic medium, subcultured (1% inoculation) the cells in this medium for five generations, and then transferred the enrichment of the fifth generation (1% inoculation) to a heterotrophic TRM medium^73^ with the following composition (L^−1^): 1 g of yeast extract, 4 g of tryptone, 23 g of NaCl, 5 g of MgCl_2_·6H_2_O, 0.7 g of KCl, 0.5 g of (NH_4_)_2_SO_4_, 3.3 g of PIPES disodium salt, 1 ml of NaBr (5%, w/v), 1 ml of SrCl_2_·6H_2_O (1%, w/v), 1 ml of KH_2_PO_4_ (5%, w/v), 1 ml of K_2_HPO_4_ (5%, w/v), 1 ml of CaCl_2_·2 H_2_O (2%, w/v), and 1 ml of Na_2_WO_4_ (10 mM). The medium was adjusted to pH 7.0, autoclaved, separated into anaerobic tubes (5 ml), and supplemented with 0.05 g of elemental sulfur and 0.05 ml of Na_2_S·9 H_2_O (10 %, w/v; pH 7.0), and the headspace of the tubes was pressurized with pure nitrogen gas. Heterotrophic enrichment was performed at 45 °C. Colonies were obtained on solid TRM medium (1.5 % Gelrite) using the rolling tube method, which was repeated three times.

### Epifluorescence microscopy and transmission electron microscopy

Fluorescent staining by SYBR Green I (Invitrogen) was carried out on cells filtered onto 0.2-μm polycarbonate GTBP membranes (Millipore) following established protocols^74^. Images were captured with a fluorescence microscope (Nikon Eclipse 90i).

For transmission electron microscopy, cells were harvested at 3,000 ×g for 5 minutes, fixed in 4% formaldehyde for 2 hours at room temperature, and washed with 1× PBS (pH 7.4). Fixed cells were negatively stained with 2 % (w/v) phosphotungstic acid (pH 6.5) for observation with a transmission electron microscope (FEI Tecnai G2 Spirit Biotwin) at an acceleration voltage of 120 kV.

### Growth characteristics

The effects of the temperature, pH, and NaCl concentration on the growth of strain 3DAC were determined in an anaerobic TRM medium with a headspace gas composed of pure nitrogen gas at atmospheric pressure supplemented with 1% (w/v) elemental sulfur. To evaluate the optimum temperature for growth, strain 3DAC was cultivated at 25-80 °C with 5 °C intervals. The ranges of pH and NaCl concentrations for growth were determined at 45 °C. The optimal pH for growth was determined by varying the pH in the culture medium between 4.0 and 9.0 using the following buffers at a concentration of 10 mM: acetate at pH 4.0, 4.5 and 5.0; MES at pH 5.5 and 6.0; PIPES at pH 6.5 and 7.0; HEPES at pH 7.5 and 8.0; and TAPS at pH 8.5 and 9.0. The pH was adjusted at room temperature before autoclaving. To test the effect of the NaCl concentration on growth, NaCl was added to the medium at final concentrations within the range of 0.5–5.0% (w/v). All of these tests were performed in triplicate in the dark without shaking. Growth was determined by monitoring the OD_600_ with a quartz colorimeter (50 mm optical path) on a spectrophotometer (HACH DR5000). The specific growth rate (*μ*; h ^−1^) was calculated according to the equation μ=ln[OD_600_(*t*2)/OD_600_(*t*1)]/(*t*2 − *t*1), where OD_600_(*t*1) and OD_600_(*t*2) are the optical densities of liquid cultures measured at 600 nm at time points *t*2 and *t*1, and *t*1 and *t*2 are the time points (in hours) of the exponential growth phases^75^. Growth rates were calculated using linear regression analysis of 4 to 6 points along the linear portions of the growth curves.

### Determination of growth pressure

The pressure range (0.1, 10, 20, 30, 40, 50, 60, 70, 80, 90 and 100 MPa) was tested with syringes loaded with TRM (2.5% NaCl, pH 7.0) supplemented with 1% (w/v) elemental sulfur. These syringes were placed into high-pressure and high-temperature reactors and incubated at 70 °C. Growth yields and growth rates were measured as previously described. All of these tests were performed in triplicate.

### Essential amino acid tests

Mother liquors of 20 kinds of amino acids (L-alanine, L-arginine, L-asparagine, L-aspartic acid, L-cysteine, L-glutamine, L-glutamic acid, glycine, L-histidine, L-isoleucine, L-leucine, L-lysine, L-methionine, L-phenylalanine, L-proline, L-serine, L-threonine, L-tryptophan, L-tyrosine, and L-valine) were prepared. Then, each combination of 19 amino acid mother liquors was added to basal TRM medium (2.5% NaCl, no yeast extract or tryptone, supplemented with 1 ml of trace element solution^76^, 10 ml of vitamin solution^77^ and 1% (w/v) elemental sulfur) to maintain each amino acid at a final concentration of 0.1 g/L. Basal TRM medium containing all 20 amino acids and no amino acids was set as the positive and negative controls, respectively. The pH of all the tested media was adjusted to 7.0 at room temperature. For all tests, positive cultures were transferred (1% inoculum) into the tested medium at least five times for confirmation of growth. All tests were performed in duplicate at atmospheric pressure and with incubation at 70 °C.

### Determination of growth requirements

Various single carbon sources were added to a basal TRM medium (2.5% NaCl, containing only 0.002% yeast extract, without any other organic carbon sources) supplemented with 1% (w/v) elemental sulfur. The following single carbon sources, all at a final concentration of 0.2% (w/v), were tested: peptone, casamino acids, beef extract, yeast extract, tryptone, sodium acetate, sodium formate, sodium citrate, sodium pyruvate, sodium oxalate, D-fructose, D-(+)-glucose, D-(+)-xylose, D-(+)-maltose, sucrose, starch, glycogen, N-acetyl-D-glucosamine, D-mannitol, L-alanine, L-arginine, L-asparagine, L-aspartic acid, L-cysteine, L-glutamine, L-glutamic acid, glycine, L-histidine, L-isoleucine, L-leucine, L-lysine, L-methionine, L-phenylalanine, L-proline, L-serine, L-threonine, L-tryptophan, L-tyrosine, and L-valine. Growth was recorded after 24 and 48 hours. Negative (uninoculated basal TRM) and positive (inoculated TRM containing yeast extract and peptone) controls were prepared for each substrate. For all tests, positive cultures were transferred (1% inoculum) into the tested medium for confirmation of growth.

To test carbon source utilization with the addition of 20 kinds of amino acids, various carbon sources at a final concentration of 0.2% (w/v) were added into a basal TRM medium (2.5% NaCl, no yeast extract or tryptone) and supplemented with 20 kinds of amino acids (each at a final concentration of 0.1 g/L) and 1% (w/v) elemental sulfur. The carbon sources used were as follows: D-(+)-glucose, D-fructose, D-(+)-maltose, sucrose, starch, sodium acetate, sodium formate, and sodium pyruvate. Basal TRM medium supplemented with 20 kinds of amino acids and 1% (w/v) elemental sulfur was used as a control.

To examine the ability of the isolate to grow in the absence of elemental sulfur, cells were cultured in TRM medium (2.5% NaCl, sodium sulfate removed) without sulfur. Other sulfur substitutes, such as sodium sulfate (20 mM), sodium sulfite (20 mM), sodium thiosulfate (20 mM), L-cysteine (20 mM) and L-methionine (20 mM), were also tested.

The pH of all the tested media was adjusted to 7.0 at room temperature. All tests were performed in duplicate at atmospheric pressure and with incubation at 70 °C. In cultures containing insoluble substrates, growth was monitored by microscopic examination.

The sulfide concentration was measured by the methylene blue method^78^ using a sulfide measurement kit (HACH) on a spectrophotometer (HACH DR5000).

### Genome sequencing and analysis

Genomic DNA was extracted by a modified SDS-based DNA extraction method^79^ and sequenced on the PacBio RSII SMRT and Illumina HiSeq X Ten sequencing platforms (BGI, China). The genome was assembled into one contig from the PacBio reads (3,668 Mbp, 2308-fold coverage) by Falcon v0.3.0^80^ and Celera Assembler v8.3^81^. Then, the assembled genome was corrected with Illumina HiSeq data (736 Mbp, 463-fold coverage) using GATK v1.6-13^82^. Protein-coding genes were predicted by Glimmer v3.02^83^, and rRNA and tRNA annotations were performed with RNAmmer v1.2^84^ and tRNAscan-SE v1.3.1^85^, respectively. Predicted gene functions were annotated with databases including NCBI-nr (update on Oct 10, 2017), KEGG (BlastKOALA, v2.2, updated on May 15, 2019)^86^, and eggNOG-Mapper v2.0.1b-2-g816e190^87^. Average amino acid identity (AAI) values were calculated by CompareM v.0.0.23 (https://github.com/dparks1134/CompareM).

### Amplicon sequencing and diversity analyses

For the chimney sample and chemoautotrophic enrichment, the bacterial V4 region of the 16S rRNA gene was amplified by B515F (5’-XXXXXXXXGTGCCAGCMGCCGCGGTAA-3’) with eight-nucleotide key tags for each sample and B806R (5’-GGACTACHVGGGTWTCTAAT-3’)^88^; the X region represents the various key tags for each sample. The PCR program was 3 min at 95 °C, followed by 35 cycles of 94 °C for 40 s, 56 °C for 1 min, and 72 °C for 1 min. The final extension step was 72 °C for 7 min. The 50 µL amplification mixture contained 1 µL of each forward and reverse primer, 1 µL template DNA, 5 µL 10× Ex Taq buffer, 0.5 µL ExTaq polymerase (TaKaRa, Japan), 4 µL of 2.5 mM dNTP mix and 37.5 µL ddwater. PCR products were purified with an E.Z.N.A. Gel Extraction Kit (OmegaBio-Tek, USA) and sequenced on the MiSeq platform (Illumina, USA) according to the manufacturer’s instructions.

Data analysis of the 16S rRNA MiSeq sequences was performed using the QIIME version 1.9.1 software pipeline^89^ and the QIIME-compatible version of the SILVA-132 database for template-based alignment and taxonomic assignment. Assembled reads that passed the chimera check were clustered into de novo operational taxonomic units (OTUs) at a cut-off of 97% sequence similarity.

### RNA sequencing and quantitative real-time PCR

Cells of strain 3DAC cultured with and without sulfur were harvested at the log phase by centrifugation. Three replicates under each condition were prepared for transcriptome analysis. Total RNA was extracted using a Total RNA Extractor (TRIzol) kit (Sangon Biotech, China) according to the manufacturer’s protocol. For RNA sequencing library preparation, genomic DNA was removed with DNase I, and ribosomal RNA (rRNA) was removed with a Ribo-off rRNA Depletion Kit (bacteria) (Vazyme Biotech Co., Ltd, China). A complementary DNA (cDNA) library was generated by a VAHTS™ Stranded mRNA-seq Library Prep Kit (Vazyme Biotech Co., Ltd, China) following the manufacturer’s instructions. Sequencing was performed on the Illumina HiSeq X Ten (Illumina, USA) platform at Sangon Biotech (Shanghai Co., Ltd., China).

The sequencing quality controls of the raw reads produced by Illumina HiSeq X Ten were assessed using FastQC^90^, and all samples passed. Low-quality reads (Q value of < 20), ambiguous “N” nucleotides, adapter sequences and fragments of < 35 bp were removed from raw reads with Trimmomatic^91^ for further analysis. rRNA was removed using a sortmerna with default settings. Read mapping of the non-rRNA to the genome of *Zhurongbacter thermophilus* 3DAC was performed using Bowtie 2^92^, and the relevant mapping information was summarized. The uniquely mapped reads were collected and analyzed with the DESeq2 package based on the Poisson distribution to identify the differentially expressed genes (DEGs). Estimation of gene expression levels was based on the transcript per million (TPM) values^93^.

To validate the RNA-Seq data, the expression levels of 10 randomly selected genes were quantified using quantitative real-time PCR (qRT-PCR). After the extraction of RNA, residual DNA was removed by a TURBO DNA-free Kit (Thermo Fisher Scientific, USA), and cDNA synthesis reverse transcription was followed by a RevertAid First Strand cDNA Synthesis Kit (Thermo Fisher Scientific, USA). The primers used for quantitative real-time PCR (RT-qPCR) were designed by Primer-BLAST^94^ (http://www.ncbi.nlm.nih.gov/tools/primer-blast) and are listed in Supplementary Table 5. Gene expression was quantified in different samples using the 16S rRNA gene as a reference gene. RT-qPCR was performed on a StepOnePlus Real-Time PCR instrument (Applied Biosystems, USA) with PowerUp SYBR Green Master Mix (Applied Biosystems, USA) using the following program: 95 °C for 10 min, followed by 40 cycles of 95 °C for 15 s and 60 °C for 1 min. All qRT-PCR analyses were performed with four technical replicates. The RNA-Seq fold changes were plotted against the qRT-PCR fold changes, and the correlation coefficients (R^2^) between these two data sets were calculated (Supplementary Fig. 2).

### Phylogenetic analyses

For the 16S rRNA gene tree, the 16S gene sequences were aligned using SINA v1.2.11^95^, and the full alignment was stripped of columns containing 70% or more gaps with a Perl script. The maximum-likelihood phylogeny of the 16S rRNA gene tree was inferred by IQ-TREE v 2.0.6^96^ with the “-MFP -B 1000” options for best-fit model selection and ultrafast bootstrap. The reference sequences of other bacteria were downloaded from the Silva database, sequences smaller than 1400 bp were filtered out, and OTUs were picked from the remaining sequences at a cutoff of 97%. The above-described representative OTU sequences were used for the tree.

For phylogenomic analysis based on conserved proteins, the representative bacterial dataset used by Coleman et al., 2021 and selected archaeal reference genomes from the different phyla were downloaded from the NCBI prokaryotic genome database (https://www.ncbi.nlm.nih.gov/assembly/). These reference genomes and the 3DAC genome from this study were used to construct a phylogenomic tree based on a concatenated alignment of five different sets of marker genes (Supplementary Table 6-9)^21, 41–43^. Specifically, each of the marker protein sequences from the reference genomes and the 3DAC genome were aligned using the MAFFT algorithm v7.313^97^ with the parameters --ep 0 --genafpair --maxiterate 1000 and filtered with trimAl v1.4.rev2^98^ with the parameter -automated1. Then, all marker protein sequences were concatenated into a single alignment, and phylogenetic trees were built using IQ-Tree v 2.0.6 with the best fitting model LG+C60+R10 with an ultrafast bootstrap value of 1000. All trees were visualized by iTOL^99^ online software.

### Evolutionary analyses of the putative superphylum Zhurongbacteria

A total of 43 cultured bacterial complete genomes (Supplementary Table 10) from the Zhurongbacteria superphylum as well as potential candidate phyla were selected for evolutionary analyses. Genomes were downloaded from NCBI genomic database and predicted with Prodigal version 2.6.3^100^. Then all genomes were annotated with eggNOG-Mapper v2.0.1b-2-g816e190^87^. The comparative genomic analysis was performed by OrthoFinder^101^ with default parameters. A total of 6,578 orthogroups from 43 representative genomes were obtained from OrthoFinder. The orthogroups that contain four or more sequences were separately aligned using the MAFFT algorithm v7.313^97^ with the parameters --ep 0 --genafpair --maxiterate 1000, filtered with trimAl v1.4.rev2^98^ with the parameter -automated1 and then trees were constructed using IQ-Tree with best-fitting model. To address the evolutionary history of the superphylum Zhurongbacteria, ancestral family gene sets were inferred using the program amalgamated likelihood estimation (ALE) version 0.4^102^. Only bacteria with complete genome information were considered in the present study because incomplete genomes would lead to biased analyses in gene gain and loss processes. Here, the phyla Synergistota and Deinococcota were used as the outgroups. The representative genomes from the phyla Aquificota, Thermosulfidibacterota and Thermodesulfobacteriota were also considered in the ALE analysis. For Zhurongbacteria, cultured bacteria with complete genomes in the phylum Dictyoglomota includes *Dictyoglomus turgidum* DSM6724 and *Dictyoglomus thermophilum* H-6-12; the phylum Thermodesulfobiota includes Thermodesulfobium acidiphilum 3127-1 GCA and Thermodesulfobium narugense DSM14796; the phylum Caldisericota includes *Caldisericum exile* AZM16c01; the phylum Zhurongbacter includes *Zhurongbacter thermophilus* 3DAC; the phylum Coprothermobacterota includes *Coprothermobacter platensis* DSM11748 and *Coprothermobacter proteolyticus* DSM5265; and the phylum Thermotogota includes *Pseudothermotoga hypogea* DSM11164, *Thermotoga neapolitana* DSM4359, *Thermosipho melanesiensis* BI429, *Fervidobacterium pennivorans* DSM9078, *Kosmotoga pacifica* SLHLJ1, *Mesotoga prima* MesG1Ag42, *Defluviitoga tunisiensis* L3, *Petrotoga mobilis* SJ95, *Marinitoga piezophila* KA3, and *Athalassotoga saccharophila* NAS-01. Ancestor metabolic reconstruction was performed with the predicted ancestral gene set (851 sequences) at the node of the superphylum Zhurongbacteria with average gene copy number larger than 0.3.

### Metabolic modeling

A genome-scale metabolic model of *Zhurongbacter thermophilus* 3DAC, GEM-i3DAC, was constructed according to the complete genome of the 3DAC strain. The draft model construction referred to ortholog mapping to public databases (KEGG^103^, EggNOG^87^, BIGG^104, 105^, ModelSeed^106^ and TCDB^107^) (performed in November 2020) and the published model of the related thermophilic bacterial strain *Thermotoga maritima* MSB8^108, 109^. Extensive manual curations were carried out following draft model reconstruction to integrate the latest biochemical evidence of enzymatic functions in (hyper)thermophiles. Overall, literature evidence was assigned to 57 reactions in the 3DAC model. The biomass equations of 3DAC were formulated based on the biosynthesis of macromolecules, including DNA, RNA, and proteins. Their biosynthesis was defined to account for the millimole composition of each building block in assembling one gram of a given component and the associated energy cost. Simulations of carbon source utilization, electron acceptors and gene deletion were formulated with exchange constraints that represent the corresponding culture conditions used in the experiments.

The KO terms of the predicted ancestral gene set were used to construct the initial genome-scale metabolic model of the last common ancestor of the superphylum Zhurongbacteria. Reactions for each KO term were obtained from the latest KEGG database (performed in March 2022). Compounds were initially introduced from the latest KEGG database, but the physiological charges of each and the modified hydrogen number of charged compounds were added. The biomass objective function was introduced from the *Escherichia coli* core model^110^, including DNA, RNA and proteins that accounted for approximately 80% of the dry weight of cells^111^ (Supplemental Data 2). Metabolic simulations were performed with exchange constraints identified based on the environmental parameters measured in typical hydrothermal vents^112^ and the potential nutrients and products revealed by metabolic reconstruction (Supplementary Data 2). Essential amino acids were identified as those missing 50% or more of the reactions from the whole biosynthesis pathway. Other amino acids were treated as nonessential. To quantitatively simulate the growth yields using the tested sole carbon sources, each carbon source was constrained to 10 mM of total carbon to avoid the influence of different carbon numbers in the formula, with all essential amino acids (1 mM for each).

For both the 3DAC model and the ancestor model, consistency checks of all reactions were computationally conducted using the *formulacheck, chargecheck* and *masscheck* functions in PSAMM version 1.1.1 (https://zhanglab.github.io/psamm/)113, and the unbalanced equations were manually curated. Gap-filling processes were performed to complete the synthesis of the main biomass components (DNA, RNA and proteins) and essential cofactors (i.e., NAD^+^/NADP^+^ and CoA) by the *fastgapfill* function in PSAMM. Flux variation analysis (FVA) to optimize biomass production was performed using the *fva* function in PSAMM with IBM ILOG CPLEX Optimizer version 12.7.1.0.

## Data and materials availability

All data are available in the main text or supplementary materials. The genome sequence of *Zhurongbacter thermophilus* 3DAC was reconstructed and deposited under accession number CP046447. Amplicon sequencing data have been deposited at SRA under accession numbers SRR14072822, SRR14072823, SRR14072817, SRR14072825, SRR14072824, SRR14072798, and SRR14072797, and transcriptome sequence data have been deposited at SRA under accession numbers SRR14072890 and SRR14072891. Biological materials are available from the corresponding author upon reasonable request.

## Acknowledgments

We thank all of the crew members of R/V *Atlantis* and ROV *Jason II* during cruise AT26-10 for their effort and help in sample collection. We thank Dr. Fengping Wang for her useful discussions in experimental design and article writing. We also thank Dr. Tom A. Williams and Edmund Moody for kindly helping with and discussing the phylogenetic analyses. This study was financially supported by the following funding: the Natural Science Foundation of China (grant numbers 41530967, 41776173, 92051116, 41902313, 20ZR1428000), the National Key R&D project of China (grant number 2018YFC0309800), as well as the US National Science Foundation grant OCE-1136727 and the *WHOI Investment in Science Fund* to S.M.S.

## Author contributions

X.X. designed and supervised the project. X.X. and S.M.S collected the chimney samples. H.L. carried out the experiments. Y.W. performed the phylogenetic and evolutionary analyses with input from H.L.. W.Z. curated the genome annotations and performed the metabolic model constructions and simulations. Y.W., L.H., and W.Z. analyzed the data. Y.W., L.H., W.Z. and X.X. wrote the manuscript. S.M.S. revised the manuscript. All authors contributed to the final version of the manuscript and approved the manuscript and supporting information.

---

**Extended Data Fig. 1.**
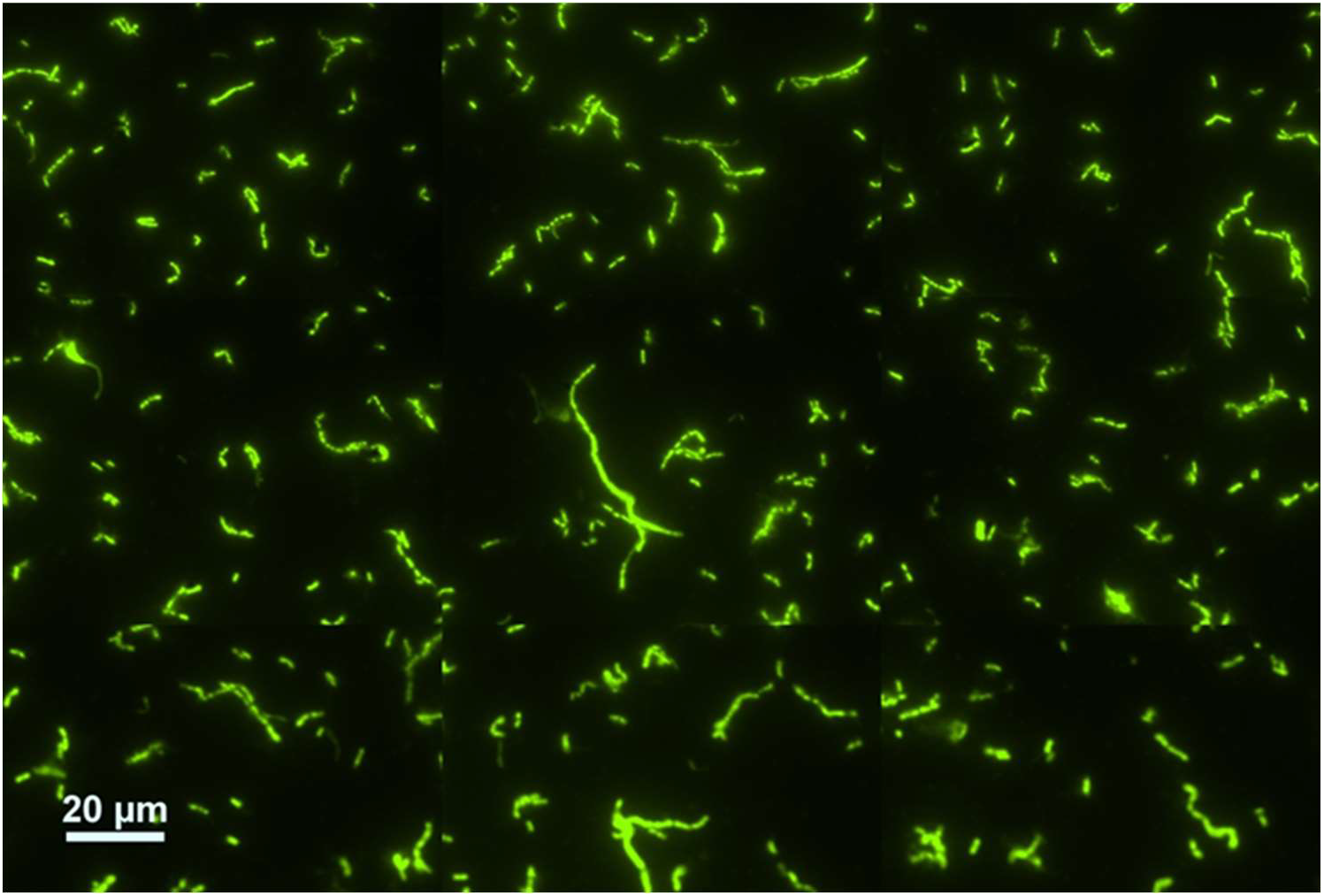
Fluorescence image of 3DAC cells stained with SYBR Green I. During growth, 3DAC cells formed long chains.

**Extended Data Fig. 2.**
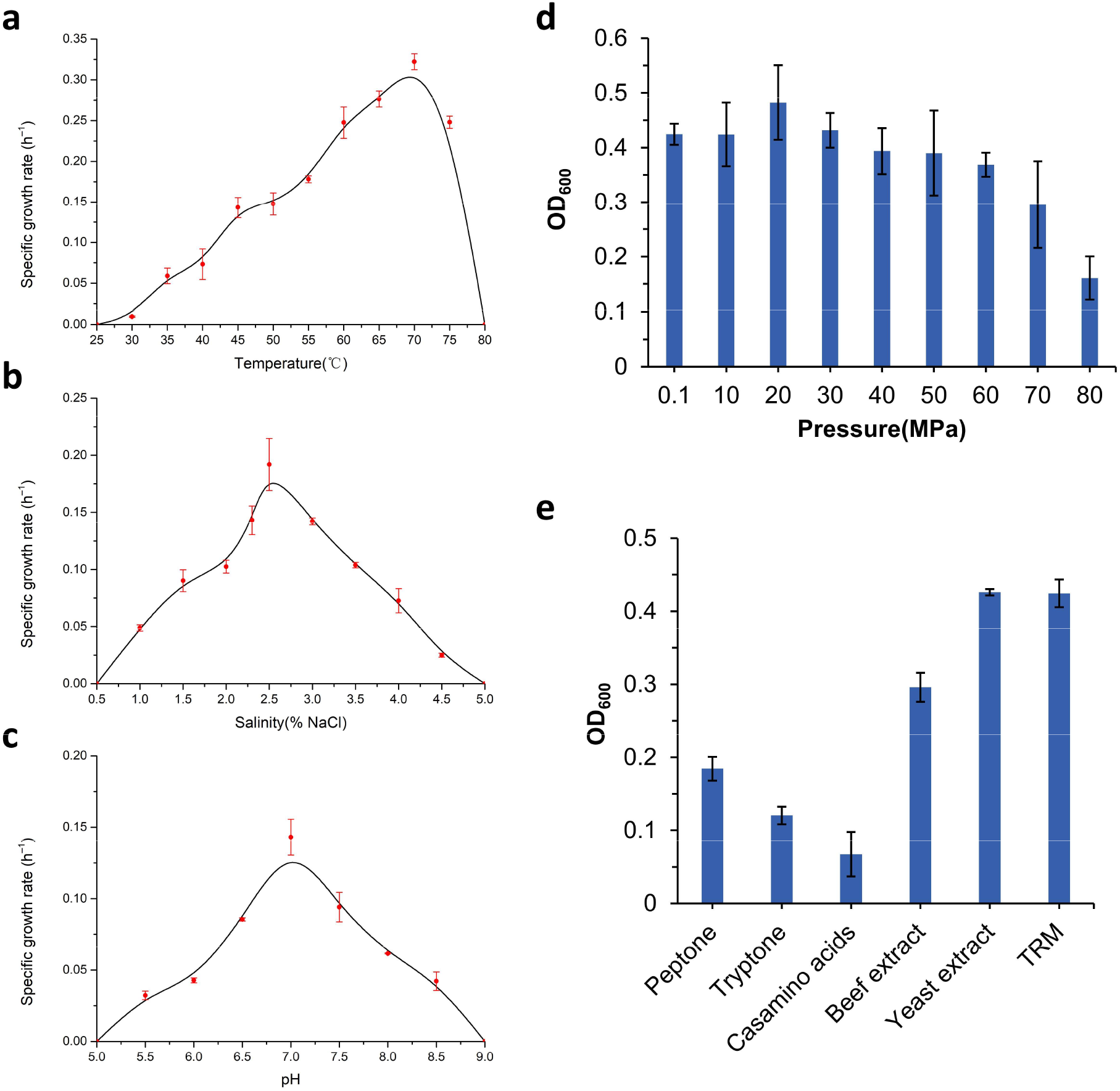
Overall characteristics of the 3DAC strain. Effects of the temperature (a), NaCl concentration (b), and pH (c) on the specific growth rate of strain 3DAC in TRM medium under atmospheric pressure. (d) Growth yields of strain 3DAC after 24 h incubations at different pressures. (e) Growth yields of strain 3DAC after 24 h incubations with different carbon sources. All the error bars in this figure represents standard deviations.

**Extended Data Fig. 3.**
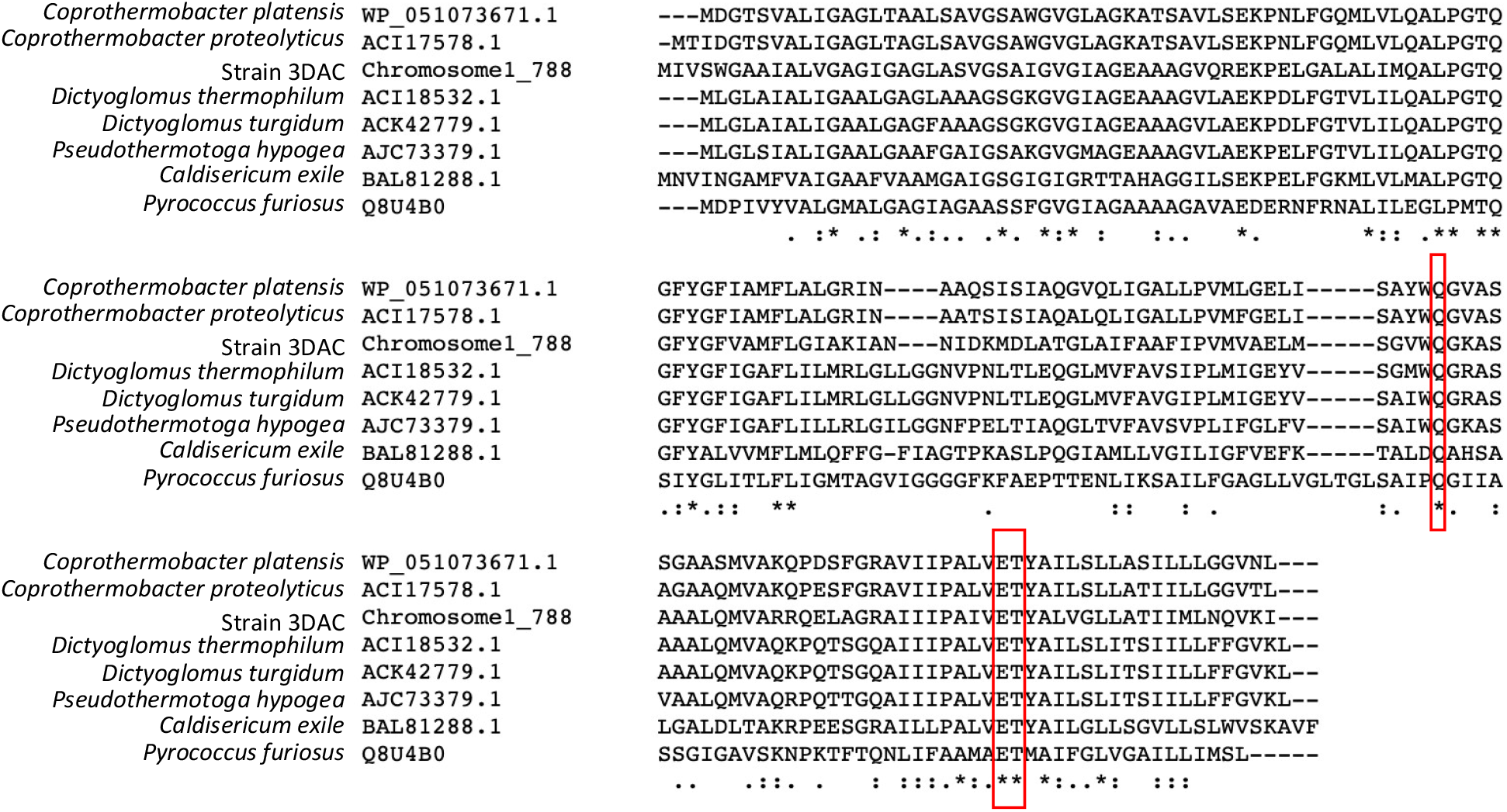
Multialignment of the c subunit in V-type ATP synthase of strain 3DAC and related species. The red boxes highlight the Na+-binding motif (“Q… ET” motif) in the c subunit ^114^. *Pyrococcus furiosus* was used as a reference. Sequence alignment was performed using Clustal Omega (Dublin, Ireland).

**Extended Data Fig. 4.**
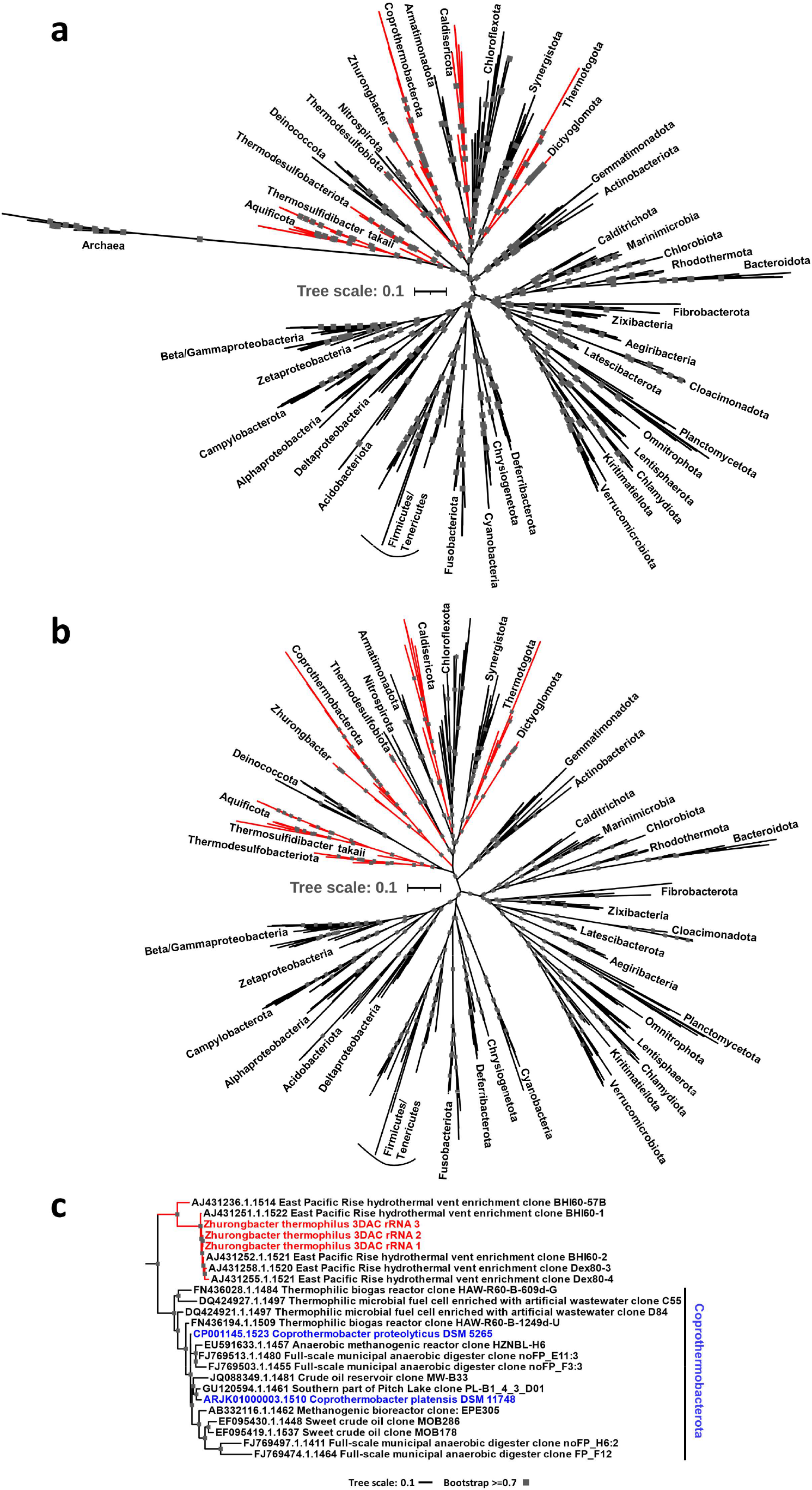
Maximum-likelihood phylogeny of the 16S rRNA gene tree. a, Tree with archaeal sequences. b, Tree without archaeal sequences. c, Detailed 16S rRNA gene sequence information of Zhurongbacterota and Coprothermobacterota. The gray squares indicate bootstrap higher than 0.7.

**Extended Data Fig. 5.**
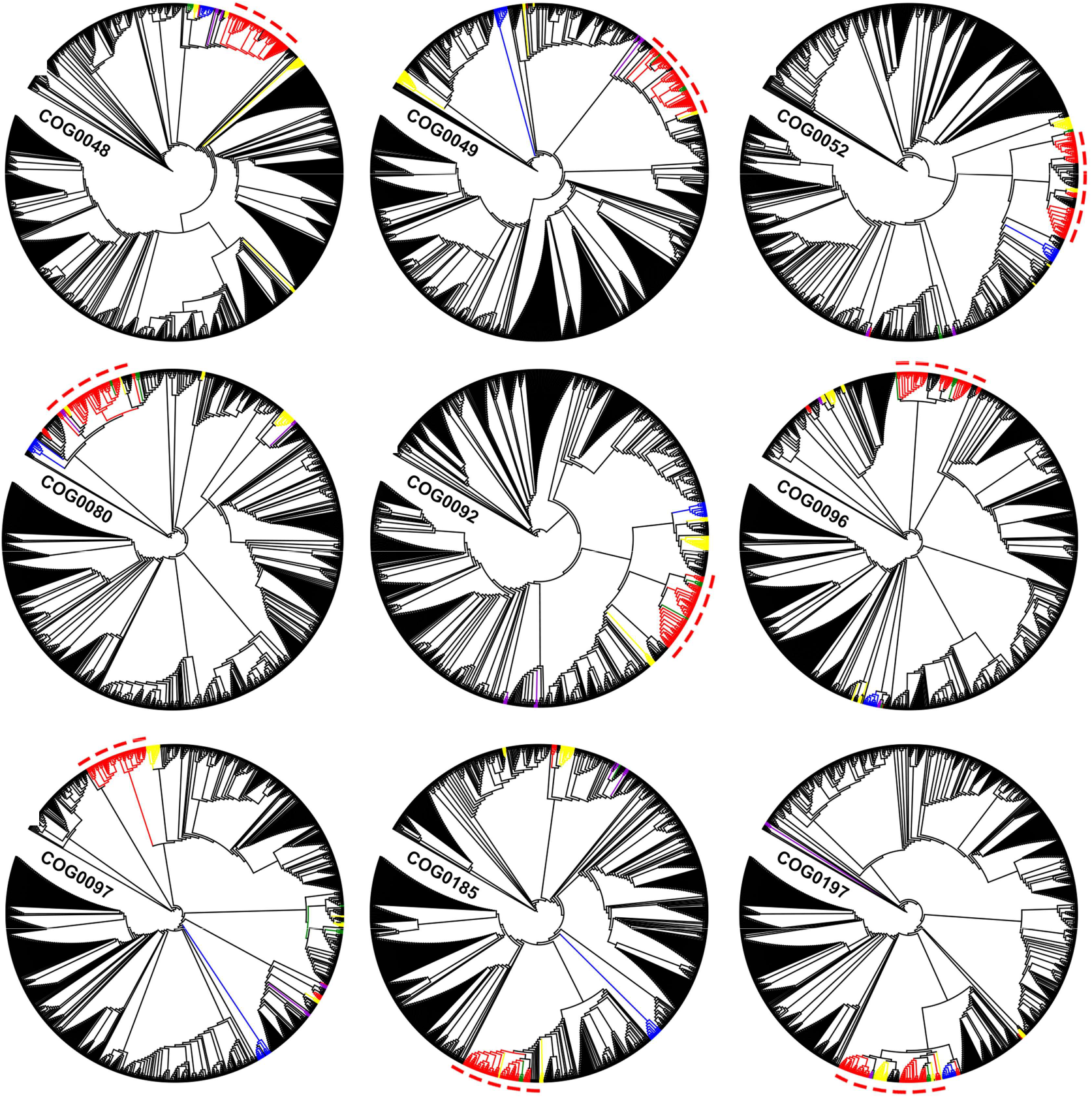
Phylogenetic trees of selected single conserved protein sequences. Red indicates the superphylum Zhurongbacteria, and Aquificota here is also colored in red. Blue represents Synergistota, while yellow represents Atribacterota and Fusobacteriota. Green indicates Caldisericota.

**Extended Data Fig. 6.**
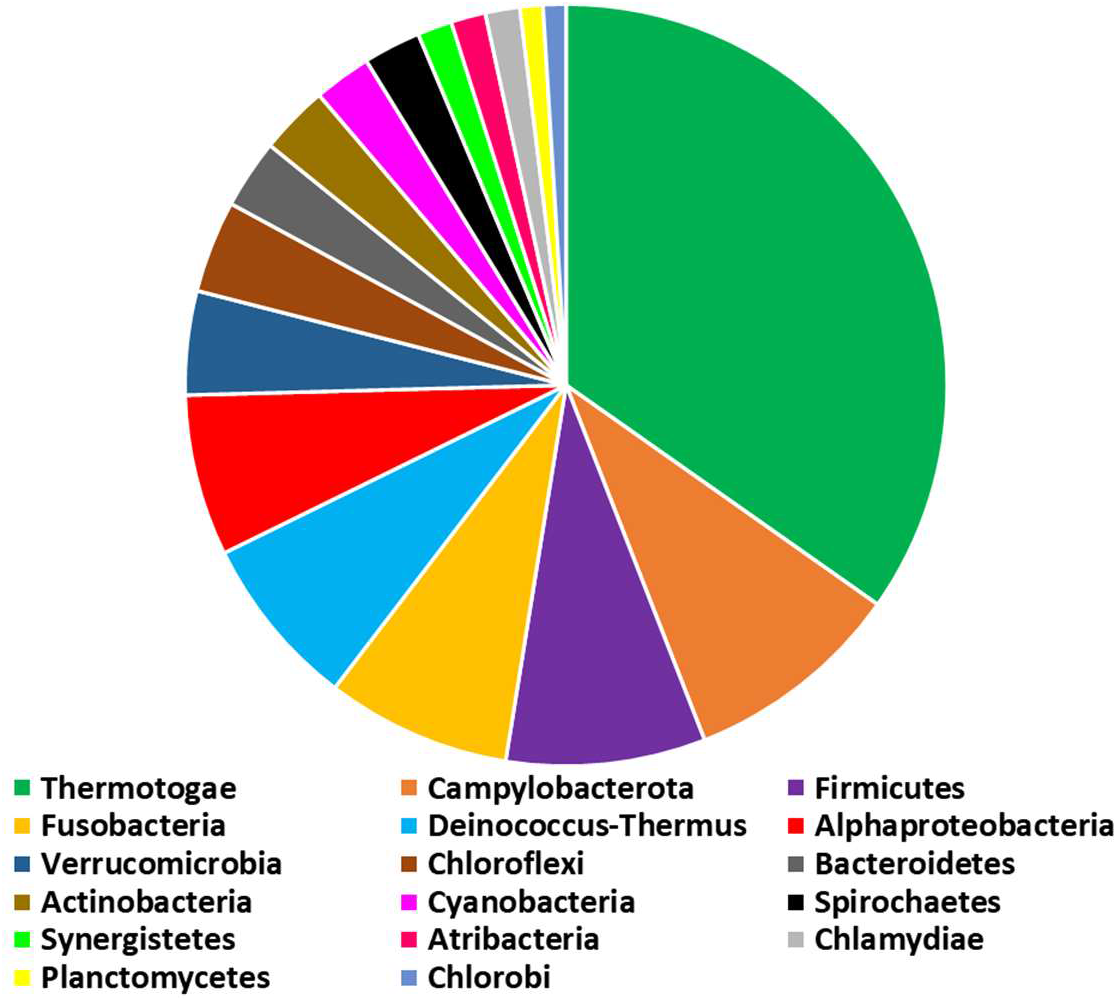
Phylogenetic results for the donor lineages for the HGTs of the conserved proteins. Thirty-seven conserved protein phylogenetic trees were constructed separately to determine whether Aquificota are closely related to the thermophilic superphylum Zhurongbacteria.

**Extended Data Fig. 7.**
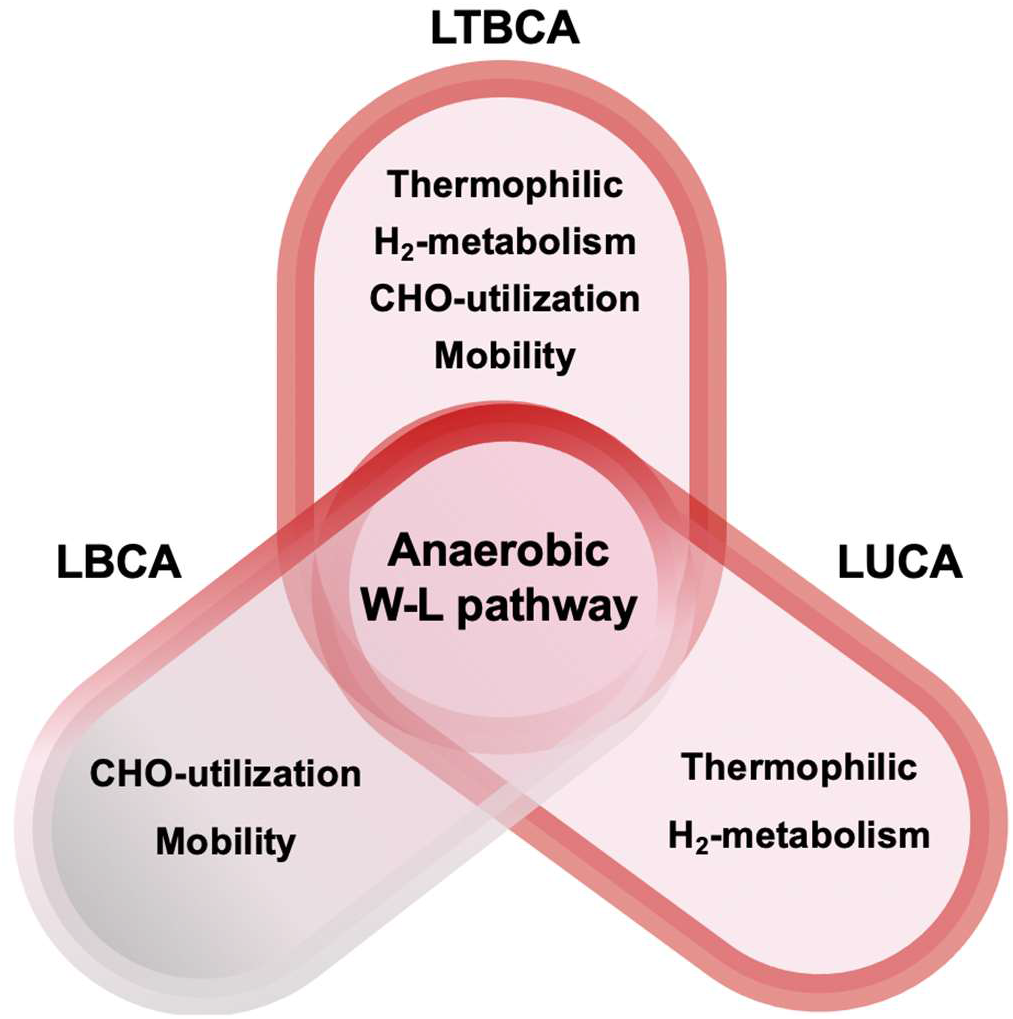
Illustration comparison of predicted metabolic features of LTBCA, LBCA and LUCA. LTBCA: last thermophilic bacterial common ancestor (This work), LBCA: last bacterial common ancestor (Coleman et al., 2021^21^, Xavier et al., 2021^67^), LUCA: last universal common ancestor (Weiss et al., 2016^17^). Anaerobic, and WL carbon fixation pathway are shared metabolic features of LTBCA, LBCA and LUCA. Besides this, LUCA shows thermophily and H2-metabolisam; LBCA shows carbohydrate utilization and mobility; LTBCA shows thermophily, H2-metabolisam, carbohydrate utilization, and mobility.

**Extended Data Fig. 8.**
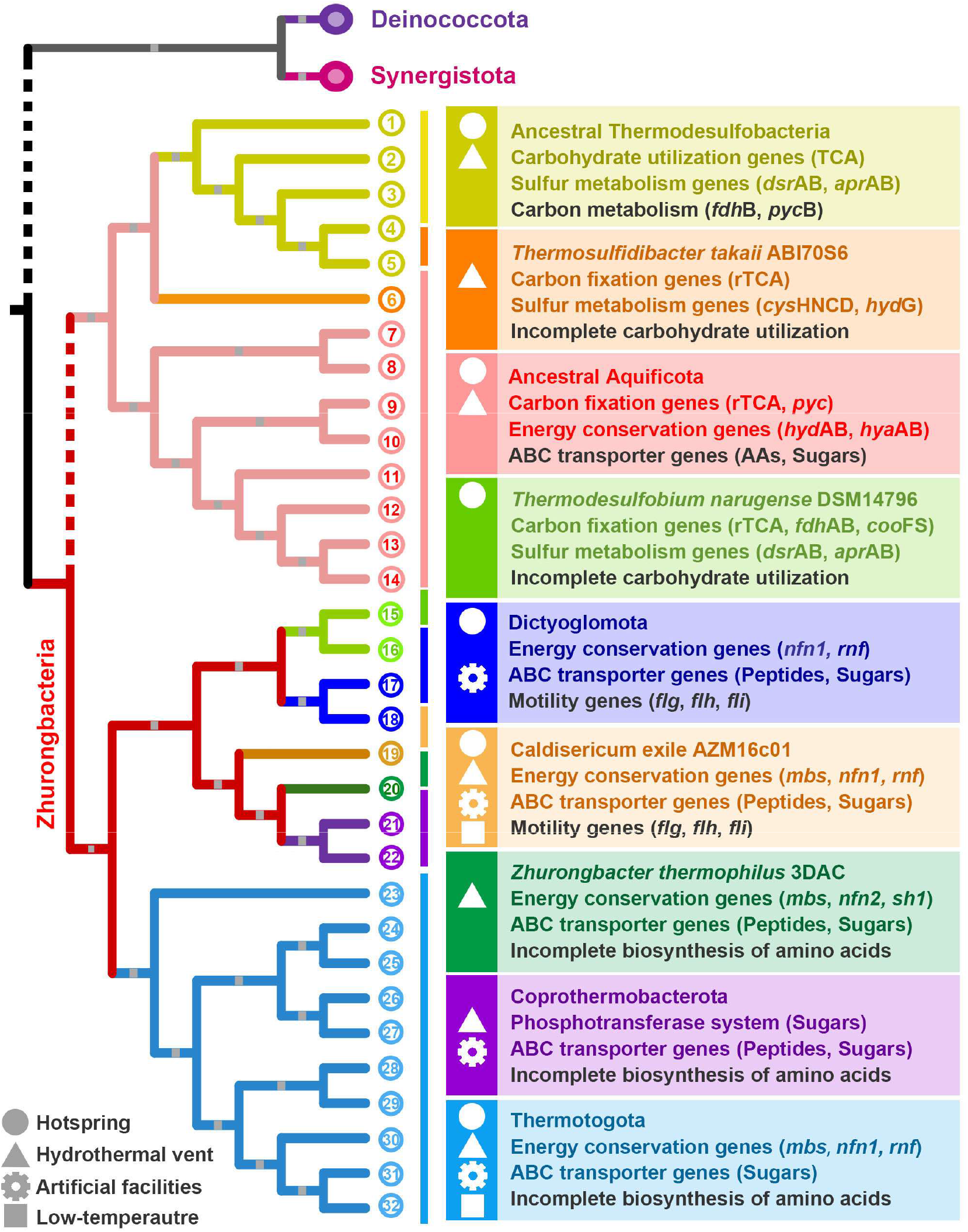
Evolutionary adaptation of the thermophilic superphylum Zhurongbacteria. Gene gain and loss of the thermophilic Zhurongbacteria superphylum. Different colors represent different phyla. Yellow: Thermodesulfobacteriota; Orange: Thermosulfidibacter; Pink: Aquificota; Grass green: Thermodesulfobiota; Blue: Dictyoglomota; Brown: Caldisericota; Dark green: Zhurongbacter; Purple: Coprothermobacterota; Cyan: Thermotogota. The complete genomes used in the current study are listed in Supplementary Table 10. The numbers and text near the ancestral branches represent the potential ancestral gene contents of each phylum.

**Extended Data Fig. 9.**
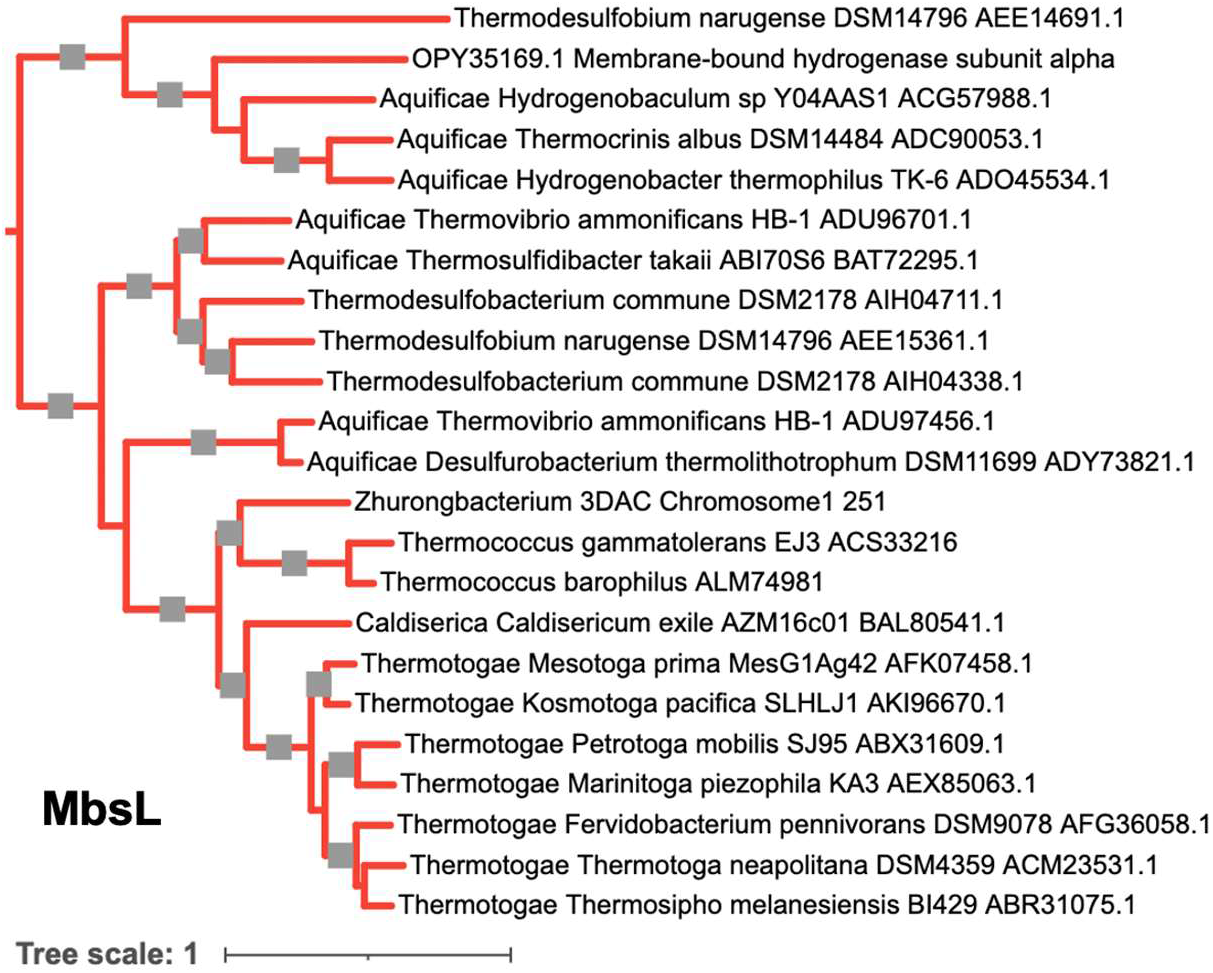
Phylogenetic tree of MbsL protein sequences from Zhurongbacteria. The gray squares indicate bootstrap higher than 0.8.

