## Supplementary for "An expanded deep-branching thermophilic bacterial clade sheds light on the early evolution of bacteria"

**This file includes the following Supplementary Information:**

Supplementary Figures 1 and 2

Supplementary Tables 1-3, 5-10

Supplementary Data descriptions

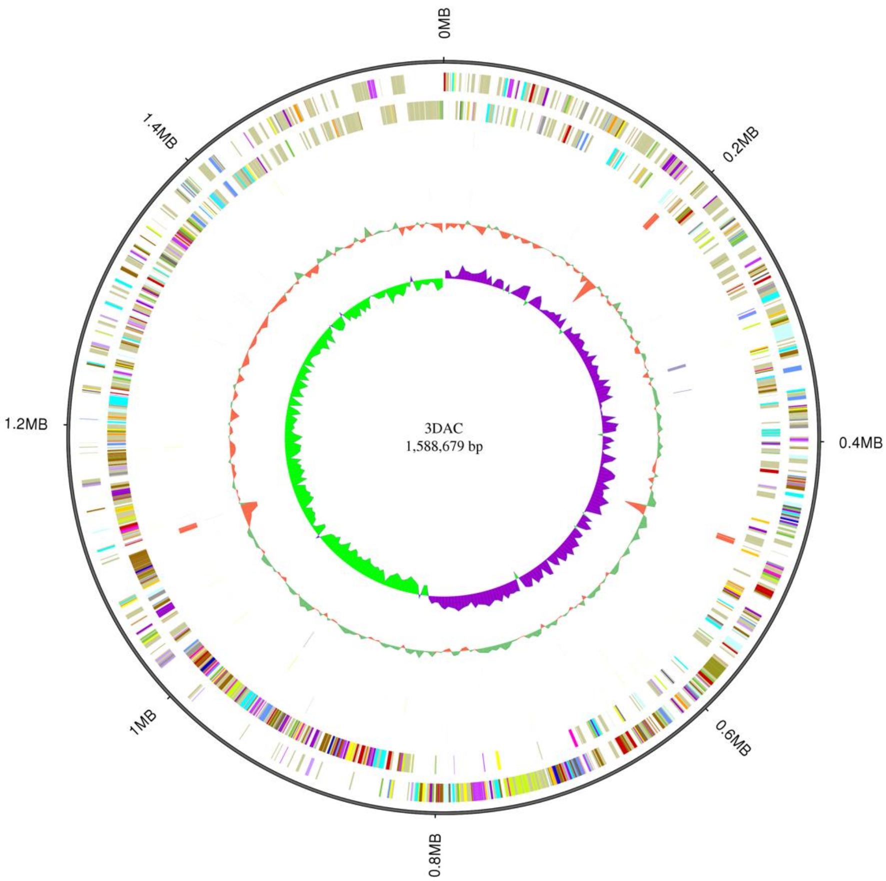


**Supplementary Figure 1 | Circular representation of the strain 3DAC genome.** Circles from the outside to the center: 1, genome size; 2, forward strand gene; 3, reverse strand gene; 4, forward strand ncRNA; 5, reverse strand ncRNA; 6, repeat; 7, GC content; 8, GC skew.

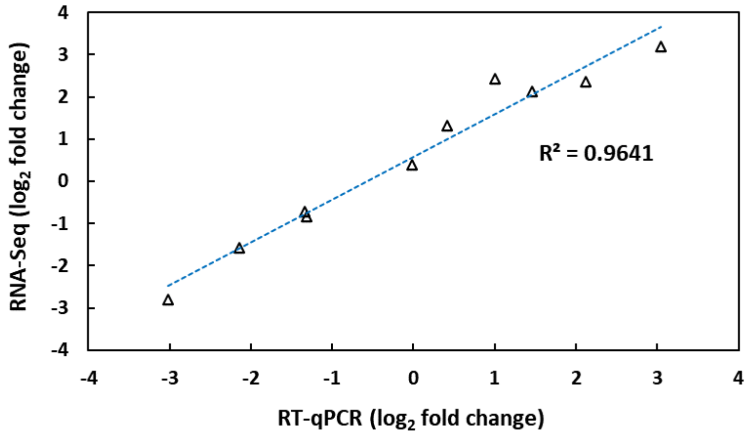


**Supplementary Figure 2 | Correlation analysis of the RNA-Seq data and RT-qPCR assays.**

The horizontal x-axis indicates the log2-fold change according to the RT-qPCR results, the vertical y-axis represents the log2-fold change according to the RNA-Seq results, and R2 indicates the R-square of the regression line.

**Supplementary Table 1.**

AAI values from strain 3DAC to close organisms calculated by compareM.

| Genome A | Genes in A | Genome B | Genes in B | Orthologous genes | Mean AAI | Std AAI | Orthologous fraction (OF) |
|---|---|---|---|---|---|---|---|
| 3DAC.Complete.genome | 1501 | Coprothermobacter_proteolyticus_SW3C_GCF_003052375.1_ASM305237v1_protein | 1371 | 524 | **49.51** | 11.62 | 34.91 |
| 3DAC.Complete.genome | 1501 | Coprothermobacter_sp_EBM-25_GCA_001896705.1_ASM189670v1_genomic | 2206 | 488 | **49.33** | 11.4 | 32.51 |
| 3DAC.Complete.genome | 1501 | Coprothermobacter_proteolyticus_BWF2A_GCF_003052365.1_ASM305236v1_protein | 1363 | 526 | **49.28** | 11.61 | 35.04 |
| 3DAC.Complete.genome | 1501 | Coprothermobacter_proteolyticus_DSM_5265_GCF_000020945.1_ASM2094v1_protein | 1408 | 560 | **49.14** | 11.62 | 37.31 |
| 3DAC.Complete.genome | 1501 | Coprothermobacter_platensis_DSM_11748_GCF_000378005.1_ASM37800v1_protein | 1400 | 554 | **48.93** | 11.37 | 36.91 |
| 3DAC.Complete.genome | 1501 | Coprothermobacter_sp_GCA_003476705.1_ASM347670v1_protein | 1257 | 424 | **48.54** | 11.55 | 28.25 |
| 3DAC.Complete.genome | 1501 | Carboxydothermus_ferrireducens_DSM11255_GCF_000427565.1_ASM42756v1_protein | 2457 | 401 | **47.13** | 10.1 | 26.72 |
| 3DAC.Complete.genome | 1501 | Thermotoga_neapolitana_DSM4359_GCF_000018945.1_ASM1894v1_protein | 1831 | 426 | **47.05** | 10.39 | 28.38 |
| 3DAC.Complete.genome | 1501 | Carboxydothermus_hydrogenoformans_Z-2901_GCF_000012865.1_ASM1286v1_protein | 2411 | 409 | **46.95** | 10.27 | 27.25 |
| 3DAC.Complete.genome | 1501 | Dictyoglomus_thermophilum_H-6-12_GCF_000020965.1_ASM2096v1_protein | 1862 | 406 | **46.76** | 10.56 | 27.05 |
| 3DAC.Complete.genome | 1501 | Thermotoga_petrophila_RKU-1_GCF_000016785.1_ASM1678v1_protein | 1780 | 435 | **46.72** | 10.52 | 28.98 |
| 3DAC.Complete.genome | 1501 | Thermoanaerobacter_thermocopriae_JCM7501_GCF_000518565.1_ASM51856v1_protein | 2297 | 408 | **46.72** | 10.11 | 27.18 |
| 3DAC.Complete.genome | 1501 | Thermotoga_naphthophila_RKU-10_GCF_000025105.1_ASM2510v1_protein | 1775 | 439 | **46.71** | 10.65 | 29.25 |
| 3DAC.Complete.genome | 1501 | Dictyoglomus_turgidum_DSM6724_GCF_000021645.1_ASM2164v1_protein | 1742 | 419 | **46.34** | 10.76 | 27.91 |
| 3DAC.Complete.genome | 1501 | Dictyoglomus_turgidum_ZAV-14_GCA_002899755.1_ASM289975v1_protein | 1633 | 354 | **46.27** | 10.72 | 23.58 |
| 3DAC.Complete.genome | 1501 | Dictyoglomus_sp_UBA8855_GCA_003509345.1_ASM350934v1_protein | 1778 | 415 | **46.2** | 10.59 | 27.65 |
| 3DAC.Complete.genome | 1501 | Ammonifex_thiophilus_GCA_003368535.1_ASM336853v1_protein | 2142 | 379 | **46.12** | 9.89 | 25.25 |
| 3DAC.Complete.genome | 1501 | Thermotoga_caldifontis_AZM44c09_GCF_000828655.1_ASM82865v1_protein | 1937 | 446 | **46.04** | 10.02 | 29.71 |

| | | | | | | | |
|---|---|---|---|---|---|---|---|
| 3DAC.Complete.genome | 1501 | Ammonifex_degensii_KC4_GCF_000024605.1_ASM2460v1_protein | 2116 | 383 | **45.98** | 10.1 | 25.52 |
| 3DAC.Complete.genome | 1501 | Acetomicrobium_mobile_DSM13181_GCF_000266925.1_ASM26692v1_protein | 1997 | 409 | **45.87** | 9.87 | 27.25 |
| 3DAC.Complete.genome | 1501 | Thermoanaerobacterium_saccharolyticum_GCF_000747665.1_ASM74766v1_protein | 2383 | 371 | **45.64** | 9.96 | 24.72 |
| 3DAC.Complete.genome | 1501 | Thermovirga_lienii_DSM17291_GCF_000233775.1_ASM23377v1_protein | 1858 | 401 | **45.59** | 9.85 | 26.72 |
| 3DAC.Complete.genome | 1501 | Acidaminococcus_intestini_RyC-MR95_GCF_000230275.1_ASM23027v1_protein | 2278 | 277 | **44.82** | 9.67 | 18.45 |
| 3DAC.Complete.genome | 1501 | Acidaminococcus_massiliensis_GCF_900095825.1_PRJEB15304_protein | 2320 | 298 | **44.62** | 9.72 | 19.85 |
| 3DAC.Complete.genome | 1501 | Dethiosulfovibrio_peptidovorans_DSM11002_GCF_000172975.1_ASM17297v1_protein | 2377 | 387 | **44.61** | 9.29 | 25.78 |
| 3DAC.Complete.genome | 1501 | Pyramidobacter_piscolens_W5455_GCF_000177335.1_ASM17733v1_protein | 2282 | 287 | **44.33** | 9.08 | 19.12 |

**Coprothermobacterota** **Thermotogota** **Dictyoglomota** **Synergistota** **Firmicutes**

**Supplementary Table 2.**

16S rRNA gene sequence identities from query sequences to top ten organisms hits by BLASTN.

**3DAC_rRNA1(1548bp)**

| Query | Organism | Query Cover | E value | Identity | Accession |
|---|---|---|---|---|---|
| 3DAC_rRNA1 | Coprothermobacter proteolyticus DSM 5265 | 99.00% | 0 | 82.88% | CP001145.1 |
| 3DAC_rRNA1 | Coprothermobacter platensis strain 3R | 95.00% | 0 | 80.97% | NR_026366.1 |
| 3DAC_rRNA1 | Fervidicola ferrireducens strain Y170 | 100.00% | 0 | 79.68% | NR_044504.1 |
| 3DAC_rRNA1 | Thermovorax subterraneus strain 70B | 96.00% | 0 | 79.49% | NR_116290.1 |
| 3DAC_rRNA1 | Thermoanaerobacter mathranii subsp. mathranii str. A3 | 100.00% | 0 | 79.18% | CP002032.1 |
| 3DAC_rRNA1 | Thermoanaerobacter thermocopriae strain JT-3 | 99.00% | 0 | 79.07% | NR_025898.1 |
| 3DAC_rRNA1 | Thermosediminibacter oceani DSM 16646 | 100.00% | 0 | 78.96% | NR_074461.1 |
| 3DAC_rRNA1 | Dictyoglomus turgidum strain DSM 6724 | 100.00% | 0 | 78.70% | NR_043385.1 |
| 3DAC_rRNA1 | Brockia lithotrophica strain Kam1851 | 96.00% | 0 | 78.70% | NR_132331.1 |
| 3DAC_rRNA1 | Dictyoglomus thermophilum H-6-12 | 100.00% | 0 | 78.49% | CP001146.1 |

**3DAC_rRNA2(1583bp)**

| Query | Organism | Query Cover | E value | Identity | Accession |
|---|---|---|---|---|---|
| 3DAC_rRNA2 | Coprothermobacter proteolyticus DSM 5265 | 99.00% | 0 | 81.25% | CP001145.1 |
| 3DAC_rRNA2 | Coprothermobacter platensis strain 3R | 95.00% | 0 | 79.42% | NR_026366.1 |
| 3DAC_rRNA2 | Thermosulfidibacter takaii ABI70S6 DNA | 100.00% | 0 | 77.11% | AP013035.1 |
| 3DAC_rRNA2 | Thermobispora bispora DSM 43833 | 100.00% | 0 | 76.04% | CP001874.1 |
| 3DAC_rRNA2 | Microbispora bispora | 100.00% | 0 | 75.85% | U83911.1 |
| 3DAC_rRNA2 | Thermosipho affectus strain ik275mar | 93.00% | 0 | 75.15% | NR_117284.1 |
| 3DAC_rRNA2 | Thermosipho globiformans strain MN14 | 92.00% | 0 | 75.07% | NR_112616.1 |
| 3DAC_rRNA2 | Sphaerobacter thermophilus DSM 20745 | 100.00% | 0 | 75.02% | CP001824.1 |
| 3DAC_rRNA2 | Thermosipho atlanticus strain DV1140 | 96.00% | 0 | 74.70% | NR_029020.1 |
| 3DAC_rRNA2 | Thermosipho melanesiensis BI429 | 100.00% | 0 | 74.54% | NR_102981.1 |

| 3DAC_rRNA3(1550bp) | | | | | |
|---|---|---|---|---|---|
| **Query** | **Organism** | **Query Cover** | **E value** | **Identity** | **Accession** |
| 3DAC_rRNA3 | Coprothermobacter proteolyticus DSM 5265 | 99.00% | 0 | 83.00% | CP001145.1 |
| 3DAC_rRNA3 | Coprothermobacter platensis strain 3R | 95.00% | 0 | 80.98% | NR_026366.1 |
| 3DAC_rRNA3 | Fervidicola ferrireducens strain Y170 | 100.00% | 0 | 79.77% | NR_044504.1 |
| 3DAC_rRNA3 | Thermovorax subterraneus strain 70B | 96.00% | 0 | 79.16% | NR_116290.1 |
| 3DAC_rRNA3 | Thermoanaerobacter mathranii subsp. mathranii str. A3 | 100.00% | 0 | 79.02% | CP002032.1 |
| 3DAC_rRNA3 | Thermosediminibacter oceani DSM 16646 | 100.00% | 0 | 78.97% | NR_074461.1 |
| 3DAC_rRNA3 | Thermoanaerobacter thermocopriae strain JT-3 | 99.00% | 0 | 78.90% | NR_025898.1 |
| 3DAC_rRNA3 | Dictyoglomus turgidum strain DSM 6724 | 100.00% | 0 | 78.63% | NR_043385.1 |
| 3DAC_rRNA3 | Thermosulfidibacter takaii ABI70S6 | 100.00% | 0 | 78.34% | AP013035.1 |
| 3DAC_rRNA3 | Thermanaeromonas toyohensis ToBE | 100.00% | 0 | 77.97% | LT838272.1 |

**Supplementary Table 3.**

16S rRNA gene sequence identities from Zhurongbacterota sequences to Coprothermobacterota sequences (>1400bp) by BLASTN.

| | | Zhurongbacterota | | | | | | | |
|---|---|---|---|---|---|---|---|---|---|
| | Sequences ID | 3DAC_rRNA1 | 3DAC_rRNA2 | 3DAC_rRNA3 | Clone_BHI60-1 | Clone_BHI60-2 | Clone_Dex80-3 | Clone_Dex80-4 | Clone_BHI60-57B |
| Coprothermobacterota | CP001145.1131856.1133378 | 0.8288 | 0.8124 | 0.828 | 0.8218 | 0.8237 | 0.8259 | 0.82 | 0.7826 |
| | CP001145.940059.941581 | 0.8288 | 0.8125 | 0.83 | 0.8232 | 0.8251 | 0.8237 | 0.8215 | 0.7839 |
| | DQ424918.1.1499 | 0.8277 | 0.8111 | 0.8268 | 0.8255 | 0.8279 | 0.8284 | 0.8238 | 0.7851 |
| | AB162803.1.1528 | 0.8271 | 0.8108 | 0.8279 | 0.823 | 0.8249 | 0.8253 | 0.8218 | 0.7839 |
| | DD224536.1.1516 | 0.8271 | 0.8107 | 0.8283 | 0.8243 | 0.8262 | 0.8248 | 0.8226 | 0.7852 |
| | DJ400146.1.1516 | 0.8271 | 0.8107 | 0.8283 | 0.8243 | 0.8262 | 0.8248 | 0.8226 | 0.7852 |
| | DQ424923.1.1502 | 0.8271 | 0.8105 | 0.8261 | 0.8246 | 0.827 | 0.8275 | 0.823 | 0.7842 |
| | JQ088349.1.1481 | 0.827 | 0.8123 | 0.8272 | 0.827 | 0.8277 | 0.8263 | 0.8236 | 0.7885 |
| | KC594816.1.1498 | 0.827 | 0.8103 | 0.8253 | 0.8245 | 0.8276 | 0.8281 | 0.8228 | 0.7831 |
| | DD224534.1.1516 | 0.8262 | 0.8099 | 0.827 | 0.823 | 0.8249 | 0.8253 | 0.8218 | 0.7839 |
| | DJ400144.1.1516 | 0.8262 | 0.8099 | 0.827 | 0.823 | 0.8249 | 0.8253 | 0.8218 | 0.7839 |
| | DD224537.1.1516 | 0.8261 | 0.8097 | 0.8276 | 0.8243 | 0.8256 | 0.8242 | 0.8208 | 0.7852 |
| | DJ400147.1.1516 | 0.8261 | 0.8097 | 0.8276 | 0.8243 | 0.8256 | 0.8242 | 0.8208 | 0.7852 |
| | KF971872.1.1401 | 0.8256 | 0.8079 | 0.8256 | 0.8244 | 0.8251 | 0.8231 | 0.8221 | 0.7933 |
| | DD224535.1.1516 | 0.8255 | 0.8091 | 0.8262 | 0.8243 | 0.8241 | 0.8237 | 0.8225 | 0.7845 |
| | DJ400145.1.1516 | 0.8255 | 0.8091 | 0.8262 | 0.8243 | 0.8241 | 0.8237 | 0.8225 | 0.7845 |
| | KM819489.1.1505 | 0.8246 | 0.808 | 0.8237 | 0.8241 | 0.8264 | 0.8256 | 0.8211 | 0.7854 |
| | FN436150.1.1500 | 0.8243 | 0.8078 | 0.8251 | 0.8243 | 0.8241 | 0.8237 | 0.8225 | 0.7845 |
| | JF808034.1.1498 | 0.8243 | 0.8077 | 0.8258 | 0.8241 | 0.824 | 0.8242 | 0.8195 | 0.7839 |
| | X69335.1.1525 | 0.8243 | 0.808 | 0.8235 | 0.8185 | 0.8204 | 0.8234 | 0.8181 | 0.78 |
| | KF208636.1.1496 | 0.8241 | 0.8076 | 0.8249 | 0.8235 | 0.8234 | 0.823 | 0.8218 | 0.7837 |
| | AB274506.1.1457 | 0.8239 | 0.8069 | 0.8247 | 0.8225 | 0.8252 | 0.8226 | 0.8183 | 0.7882 |
| | FN436187.1.1499 | 0.823 | 0.8064 | 0.8229 | 0.8243 | 0.8253 | 0.8244 | 0.8225 | 0.7833 |
| | AB233998.1.1458 | 0.8225 | 0.8055 | 0.8212 | 0.8198 | 0.8203 | 0.8212 | 0.82 | 0.7864 |
| | AB537980.1.1455 | 0.8225 | 0.8055 | 0.8233 | 0.821 | 0.8217 | 0.8211 | 0.8185 | 0.7871 |
| | DQ424920.1.1497 | 0.8224 | 0.8059 | 0.8232 | 0.8219 | 0.8238 | 0.8242 | 0.8207 | 0.7828 |
| | EU812974.1.1456 | 0.8222 | 0.8051 | 0.8215 | 0.8186 | 0.8207 | 0.8223 | 0.8182 | 0.788 |

| | | | | | | | | |
|---|---|---|---|---|---|---|---|---|
| FN436144.1.1500 | 0.822 | 0.8056 | 0.8228 | 0.8215 | 0.8228 | 0.823 | 0.82 | 0.7828 |
| FN436126.1.1501 | 0.8218 | 0.8054 | 0.8226 | 0.824 | 0.8238 | 0.8234 | 0.8222 | 0.7854 |
| AB794885.1.1459 | 0.8206 | 0.8037 | 0.8214 | 0.8199 | 0.8198 | 0.8193 | 0.8181 | 0.7878 |
| AB742088.1.1458 | 0.8198 | 0.8029 | 0.8206 | 0.8184 | 0.8189 | 0.8193 | 0.8167 | 0.7857 |
| FN436177.1.1500 | 0.8197 | 0.8033 | 0.8205 | 0.8205 | 0.8218 | 0.8226 | 0.8187 | 0.7815 |
| AB630185.1.1461 | 0.819 | 0.8022 | 0.8198 | 0.8187 | 0.8189 | 0.8198 | 0.8168 | 0.7864 |
| HV515135.1.1459 | 0.8188 | 0.802 | 0.8196 | 0.8184 | 0.8187 | 0.8196 | 0.8166 | 0.7864 |
| HW083398.1.1459 | 0.8188 | 0.802 | 0.8196 | 0.8184 | 0.8187 | 0.8196 | 0.8166 | 0.7864 |
| ARJK01000003.207993.209502 | 0.8174 | 0.8024 | 0.8174 | 0.8126 | 0.8163 | 0.816 | 0.811 | 0.7802 |
| ARJK01000003.394918.396427 | 0.8174 | 0.8024 | 0.8174 | 0.8126 | 0.8163 | 0.816 | 0.811 | 0.7802 |
| GU120594.1.1461 | 0.8161 | 0.8005 | 0.8147 | 0.8139 | 0.8137 | 0.812 | 0.8095 | 0.7753 |
| FJ769513.1.1480 | 0.8128 | 0.7973 | 0.8115 | 0.8074 | 0.809 | 0.8087 | 0.8063 | 0.7772 |
| Y08935.1.1446 | 0.8097 | 0.7942 | 0.8098 | 0.8083 | 0.8106 | 0.8103 | 0.8051 | 0.7829 |
| FJ769497.1.1411 | 0.8084 | 0.7912 | 0.8074 | 0.8054 | 0.8046 | 0.8065 | 0.8068 | 0.7785 |
| FJ769503.1.1455 | 0.8064 | 0.7909 | 0.805 | 0.8037 | 0.8053 | 0.8042 | 0.8024 | 0.7743 |
| EF095435.1.1534 | 0.8057 | 0.7903 | 0.8048 | 0.8011 | 0.8048 | 0.8039 | 0.7977 | 0.7684 |
| FN436194.1.1509 | 0.8048 | 0.7888 | 0.8056 | 0.807 | 0.8068 | 0.8175 | 0.8162 | 0.7901 |
| EU591633.1.1457 | 0.7996 | 0.7836 | 0.8003 | 0.7982 | 0.7995 | 0.7991 | 0.7961 | 0.7836 |
| DQ424921.1.1497 | 0.7944 | 0.7787 | 0.7952 | 0.7942 | 0.794 | 0.7917 | 0.7976 | 0.8122 |
| FN436028.1.1484 | 0.7802 | 0.7647 | 0.781 | 0.7822 | 0.7797 | 0.7835 | 0.7829 | 0.8018 |
| DQ424927.1.1497 | 0.7747 | 0.7595 | 0.7758 | 0.7865 | 0.7857 | 0.7891 | 0.7883 | 0.8239 |
| FJ769474.1.1464 | 0.7704 | 0.7573 | 0.7693 | 0.767 | 0.7672 | 0.7693 | 0.7695 | 0.7577 |
| **Identity range** | **0.7704-0.8288** | **0.7573-0.8124** | **0.7693-0.83** | **0.767-0.827** | **0.7672-0.8279** | **0.7693-0.8284** | **0.7695-0.8238** | **0.7577-0.8239** |

**Supplementary Table 5.**

Primers used for RT-qPCR.

| Primer | Primer sequence (5'-3') |
|---|---|
| 16-F | GGATGCCCTACCACTTCAGG |
| 16-R | CACCTCGCTCTGGTGGATTT |
| 71-F | GCCCCATAGGTTCACTGCTT |
| 71-R | GTATGAAGGGCACCACACCA |
| 235-F | CTGGCATTACCGTGGGTGTA |
| 235-R | GCTCCAGTTTCAGGGCATCT |
| 246-F | AAAGCACTTGGACTGGGTGT |
| 246-R | CCACGAGGTCACCATCTCAA |
| 663-F | AAGGTTGCCTCTAAAGGGGTT |
| 663-R | TTCGGCCTTTGCCAACATAC |
| 749-F | TGGGAAGACCGTTTGTTGGG |
| 749-R | TACCAGGCGGACGACATTTT |
| 1017-F | TGCCGATGAGCTGAAGGAAA |
| 1017-R | GGTAATCTCCTCACCGCCAA |
| 1173-F | TTGCTGTGGTCTCCTTGCTT |
| 1173-R | TCTGATTGGCAGGCAGTGTT |
| 1377-F | AGCCGCAGAAGAAAGCATCT |
| 1377-R | AACACCCTAACCAGCCATCC |
| 1388-F | TTTCCCCGTAATGGCAGGAG |
| 1388-R | AATGATAAGCACCGCCGTCA |
| 16S rRNA-F | CACGGGAAACCGTGGGTAAT |
| 16S rRNA-R | GTGAGCCGTTACCTCACCAA |

**Supplementary Table 6.**

The 16 conserved protein sequences information, * are selected for eight conserved protein sequences phylogenetic tree.

| COG ID | Gene name |
|---|---|
| COG0051 | Ribosomal protein S10 |
| COG0087* | Ribosomal protein L3 |
| COG0088 | Ribosomal protein L4 |
| COG0090 | Ribosomal protein L2 |
| COG0091 | Ribosomal protein L22 |
| COG0092 | Ribosomal protein S3 |
| COG0093 | Ribosomal protein L14 |
| COG0094* | Ribosomal protein L5 |
| COG0096* | Ribosomal protein S8 |
| COG0097* | Ribosomal protein L6P/L9E |
| COG0185* | Ribosomal protein S19 |
| COG0186* | Ribosomal protein S17 |
| COG0197 | Ribosomal protein L16/L10AE |
| COG0198 | Ribosomal protein L24 |
| COG0200* | Ribosomal protein L15 |
| COG0256* | Ribosomal protein L18 |

**Supplementary Table 7.**

37 conserved protein sequences information.

| COG ID | Gene name |
|---|---|
| COG0016 | Phenylalanyl-tRNA synthetase alpha subunit |
| COG0048 | Ribosomal protein S12 |
| COG0049 | Ribosomal protein S7 |
| COG0051 | Ribosomal protein S10 |
| COG0052 | Ribosomal protein S2 |
| COG0072 | Phenylalanyl-tRNA synthetase beta subunit |
| COG0080 | Ribosomal protein L11 |
| COG0081 | Ribosomal protein L1 |
| COG0087 | Ribosomal protein L3 |
| COG0088 | Ribosomal protein L4 |
| COG0090 | Ribosomal protein L2 |
| COG0091 | Ribosomal protein L22 |
| COG0092 | Ribosomal protein S3 |
| COG0093 | Ribosomal protein L14 |
| COG0094 | Ribosomal protein L5 |
| COG0096 | Ribosomal protein S8 |
| COG0097 | Ribosomal protein L6P/L9E |
| COG0098 | Ribosomal protein S5 |
| COG0099 | Ribosomal protein S13 |
| COG0100 | Ribosomal protein S11 |
| COG0103 | Ribosomal protein S9 |
| COG0127 | Xanthosine triphosphate pyrophosphatase |
| COG0149 | Triosephosphate isomerase |
| COG0164 | Ribonuclease HII |
| COG0184 | Ribosomal protein S15P/S13E |
| COG0185 | Ribosomal protein S19 |
| COG0186 | Ribosomal protein S17 |
| COG0197 | Ribosomal protein L16/L10E |
| COG0198 | Ribosomal protein L24 |
| COG0200 | Ribosomal protein L15 |
| COG0244 | Ribosomal protein L10 |
| COG0256 | Ribosomal protein L18 |
| COG0343 | Queuine/archaeosine tRNA-ribosyltransferase |
| COG0504 | CTP synthase (UTP-ammonia lyase) |
| COG0532 | Translation initiation factor 2 (IF-2; GTPase) |
| COG0533 | Metal-dependent proteases with possible chaperone activity |
| COG0541 | Signal recognition particle GTPase |

**Supplementary Table 8.**

57 conserved protein sequences information.

| COG ID | Gene name |
|---|---|
| COG0013 | Alanyl-tRNA synthetase |
| COG0016 | Phenylalanyl-tRNA synthetase alpha subunit |
| COG0018 | Arginyl-tRNA synthetase |
| COG0048 | Ribosomal protein S12 |
| COG0049 | Ribosomal protein S7 |
| COG0051 | Ribosomal protein S10 |
| COG0052 | Ribosomal protein S2 |
| COG0060 | Isoleucyl-tRNA synthetase |
| COG0072 | Phenylalanyl-tRNA synthetase beta subunit |
| COG0080 | Ribosomal protein L11 |
| COG0081 | Ribosomal protein L1 |
| COG0085 | DNA-directed RNA polymerase beta subunit (RpoB) |
| COG0086 | DNA-directed RNA polymerase beta prime subunit (RpoC) |
| COG0087 | Ribosomal protein L3 |
| COG0088 | Ribosomal protein L4 |
| COG0089 | Ribosomal protein L23 |
| COG0090 | Ribosomal protein L2 |
| COG0091 | Ribosomal protein L22 |
| COG0092 | Ribosomal protein S3 |
| COG0093 | Ribosomal protein L14 |
| COG0094 | Ribosomal protein L5 |
| COG0096 | Ribosomal protein S8 |
| COG0097 | Ribosomal protein L6P |
| COG0098 | Ribosomal protein S5 |
| COG0099 | Ribosomal protein S13 |
| COG0100 | Ribosomal protein S11 |
| COG0102 | Ribosomal protein L13 |
| COG0103 | Ribosomal protein S9 |
| COG0127 | Xanthosine triphosphate pyrophosphatase |
| COG0130 | Pseudouridine synthase |
| COG0149 | Triosephosphate isomerase |
| COG0164 | Ribonuclease HII |
| COG0172 | Seryl-tRNA synthetase |
| COG0184 | Ribosomal protein S15P |
| COG0185 | Ribosomal protein S19 |
| COG0186 | Ribosomal protein S17 |
| COG0193 | Peptidyl-tRNA hydrolase |
| COG0197 | Ribosomal protein L16 |

| | |
|---|---|
| COG0198 | Ribosomal protein L24 |
| COG0200 | Ribosomal protein L15 |
| COG0201 | Preprotein translocase subunit SecY |
| COG0202 | DNA-directed RNA polymerase alpha subunit (RpoA) |
| COG0216 | Protein chain release factor A |
| COG0233 | Ribosome recycling factor |
| COG0244 | Ribosomal protein L10 |
| COG0255 | Ribosomal protein L29 |
| COG0256 | Ribosomal protein L18 |
| COG0343 | Queuine/archaeosine tRNA-ribosyltransferase |
| COG0481 | Membrane GTPase LepA |
| COG0495 | Leucyl-tRNA synthetase |
| COG0504 | CTP synthase |
| COG0519 | GMP synthase PP-ATPase domain |
| COG0532 | Translation initiation factor 2 |
| COG0533 | Metal-dependent proteases with possible chaperone activity |
| COG0541 | Signal recognition particle GTPase |
| COG0691 | TmRNA-binding protein |
| COG0858 | Ribosome-binding factor A |

**Supplementary Table 9.**

62 conserved protein sequences information.

| KO number | Gene name | Annotation |
|---|---|---|
| K06942 | ychF | Redox Regulated ATPase YchF |
| K01872 | AARS, alaS | Alanine tRNA ligase |
| K01889 | FARSA, pheS | Phenylalanine--tRNA ligase alpha subunit |
| K02992 | RP-S7, MRPS7, rpsG | 30S ribosomal protein S7 |
| K02358 | tuf, TUFM | Elongation factor Tu |
| K02946 | RP-S10, MRPS10, rpsJ | 30S ribosomal protein S10 |
| K02967 | RP-S2, MRPS2, rpsB | 30S ribosomal protein S2 |
| K02112 | ATPF1B, atpD | F-type H+/Na+-transporting ATPase subunit beta |
| K02867 | RP-L11, MRPL11, rplK | 50S ribosomal protein L11 |
| K02863 | RP-L1, MRPL1, rplA | 50S ribosomal protein L1 |
| K03043 | rpoB | DNA-directed RNA polymerase subunit beta |
| K03046 | rpoC | DNA-directed RNA polymerase subunit beta' |
| K02906 | RP-L3, MRPL3, rplC | 50S ribosomal protein L3 |
| K02886 | RP-L2, MRPL2, rplB | 50S ribosomal protein L2 |
| K02982 | RP-S3, rpsC | 30S ribosomal protein S3 |
| K02874 | RP-L14, MRPL14, rplN | 50S ribosomal protein L14 |
| K02931 | RP-L5, MRPL5, rplE | 50S ribosomal protein L5 |
| K02994 | RP-S8, rpsH | 30S ribosomal protein S8 |
| K02933 | RP-L6, MRPL6, rplF | 50S ribosomal protein L6 |
| K02988 | RP-S5, MRPS5, rpsE | 30S ribosomal protein S5 |
| K02952 | RP-S13, rpsM | 30S ribosomal protein S13 |
| K02948 | RP-S11, MRPS11, rpsK | 30S ribosomal protein S11 |
| K02871 | RP-L13, MRPL13, rplM | 50S ribosomal protein L13 |
| K02996 | RP-S9, MRPS9, rpsI | 30S ribosomal protein S9 |
| K00927 | PGK, pgk | Phosphoglycerate kinase |
| K01803 | TPI, tpiA | Triose phosphate isomerase |
| K02433 | gatA, QRSL1 | aspartyl-tRNA(Asn)/glutamyl-tRNA(Gln) amidotransferase subunit A |
| K03470 | rnhB | Ribonuclease HII |
| K02956 | RP-S15, MRPS15, rpsO | 30S ribosomal protein S15 |
| K02965 | RP-S19, rpsS | 30S ribosomal protein S19 |
| K02470 | gyrB | DNA gyrase subunit B |
| K02469 | gyrA | DNA gyrase subunit A |

| | | |
|---|---|---|
| K02878 | RP-L16, MRPL16, rplP | 50S ribosomal protein L16 |
| K03076 | secY | Protein translocase subunit SecY |
| K02835 | prfA, MTRF1, MRF1 | Peptide chain release factor 1 |
| K02935 | RP-L7, MRPL12, rplL | 50S ribosomal protein L7/12 |
| K02838 | frr, MRRF, RRF | Ribosome recycling factor |
| K02601 | nusG | Transcription termination antitermination protein NusG |
| K01972 | E6.5.1.2, ligA, ligB | DNA ligase NAD |
| K03438 | mraW, rsmH | 16S rRNA (cytosine1402-N4)-methyltransferase |
| K02887 | RP-L20, MRPL20, rplT | 50S ribosomal protein L20 |
| K02314 | dnaB | Replicative DNA helicase |
| K00554 | trmD | tRNA (guanine37-N1)-methyltransferase |
| K06187 | recR | Recombination protein RecR |
| K02518 | infA | Translation initiation factor IF 1 |
| K04077 | groEL, HSPD1 | Molecular chaperone GroEL |
| K03553 | recA | DNA recombination repair protein RecA |
| K02355 | fusA, GFM, EFG | Translation elongation factor G |
| K01873 | VARS, valS | Valine--tRNA ligase |
| K03702 | uvrB | Excinuclease ABC subunit B |
| K03685 | rnc, DROSHA, RNT1 | Ribonuclease III |
| K02337 | dnaE | DNA polymerase III subunit alpha |
| K02313 | dnaA | Chromosomal replication initiator protein |
| K03070 | secA | Protein translocase subunit SecA |
| K03664 | smpB | SsrA binding protein |
| K01358 | clpP, CLPP | ATP dependent Clp protease proteolytic subunit |
| K02335 | polA | DNA polymerase I |
| K03590 | ftsA | Cell division protein FtsA |
| K04485 | radA, sms | DNA repair protein RadA/Sms |
| K00962 | pnp, PNPT1 | Polyribonucleotide nucleotidyltransferase |
| K15034 | yaeJ | Aminoacyl tRNA hydrolase, ribosome associated protein |
| K03551 | ruvB | Holliday junction branch migration DNA helicase RuvB |

**Supplementary Table 10.**

Complete genomes used in ancestral reconstruction. Genome completeness and contamination were determined by CheckM.

| Lineage | Organism | Accession | Completeness | Contamination |
|---|---|---|---|---|
| Deinococcota | Truepera radiovictrix DSM17093 | GCA_000092425.1 | 97.88 | 0.42 |
| Deinococcota | Deinococcus radiophilus ATCC27603 | GCA_020889625.1 | 94.49 | 0.64 |
| Deinococcota | Thermus thermophilus HB8 | GCA_000091545.1 | 99.58 | 0 |
| Deinococcota | Meiothermus silvanus DSM 9946 | GCA_000092125.1 | 99.79 | 0.14 |
| Synergistota | Jonquetella anthropi DSM 22815 | GCA_000237805.1 | 100 | 0 |
| Synergistota | Thermovirga lienii DSM 17291 | GCA_000233775.1 | 100 | 0 |
| Synergistota | Acetomicrobium mobile DSM 13181 | GCA_000266925.1 | 100 | 0 |
| Synergistota | Aminobacterium colombiense DSM 12261 | GCA_000025885.1 | 100 | 0 |
| Synergistota | Cloacibacillus porcorum CL-84 | GCA_001701045.1 | 100 | 0 |
| Synergistota | Aminomonas paucivorans DSM 12260 | GCA_000165795.1 | 98.31 | 0 |
| Synergistota | Thermanaerovibrio velox DSM 12556 | GCA_000237825.1 | 100 | 0 |
| Thermodesulfobacteriota | Thermosulfuriphilus ammonigenes ST65 | GCA_011207455.1 | 99.59 | 1.63 |
| Thermodesulfobacteriota | Thermodesulfatator indicus DSM 15286 | GCA_000217795.1 | 99.04 | 0 |
| Thermodesulfobacteriota | Thermosulfurimonas marina SU872 | GCA_012317585.1 | 99.04 | 0.41 |
| Thermodesulfobacteriota | Caldimicrobium thiodismutans TF1 | GCA_001548275.1 | 97.94 | 1.13 |
| Thermodesulfobacteriota | Thermodesulfobacterium commune DSM 2178 | GCA_000734015.1 | 99.17 | 0 |
| Thermosulfidibacterota | Thermosulfidibacter takaii ABI70S6 | GCA_001547735.1 | 98.17 | 1.63 |
| Aquificota | Thermovibrio ammonificans HB-1 | GCA_000185805.1 | 99.58 | 0.84 |
| Aquificota | Desulfurobacterium thermolithotrophum DSM 11699 | GCA_000191045.1 | 99.16 | 1.69 |
| Aquificota | Sulfurihydrogenibium azorense Az-Fu1 | GCA_000021545.1 | 99.39 | 0 |
| Aquificota | Persephonella marina EX-H1 | GCA_000021565.1 | 99.8 | 1.83 |
| Aquificota | Hydrogenobaculum sp. HO | GCA_000341855.1 | 99.59 | 0 |
| Aquificota | Aquifex aeolicus VF5 | GCA_000008625.1 | 98.98 | 1.22 |
| Aquificota | Hydrogenobacter thermophilus TK-6 | GCA_000010785.1 | 98.78 | 0.51 |
| Aquificota | Thermocrinis albus DSM 14484 | GCA_000025605.1 | 99.39 | 0 |
| Thermodesulfobiota | Thermodesulfobium narugense DSM 14796 | GCA_000212395.1 | 98.28 | 1.72 |
| Thermodesulfobiota | Thermodesulfobium acidiphilum 3127-1 | GCA_003057965.1 | 98.28 | 1.72 |
| Dictyoglomota | Dictyoglomus thermophilum H-6-12 | GCA_000020965.1 | 100 | 1.72 |
| Dictyoglomota | Dictyoglomus turgidum DSM 6724 | GCA_000021645.1 | 100 | 0 |
| Caldisericota | Caldisericum exile AZM16c01 | GCA_000284335.1 | 98.21 | 0 |
| Zhurongbacter | Zhurongbacter thermophilus 3DAC | CP046447 | 98.21 | 0 |
| Coprothermobacterota | Coprothermobacter platensis DSM 11748 | GCA_000378005.1 | 98.21 | 0 |
| Coprothermobacterota | Coprothermobacter proteolyticus DSM 5265 | GCA_000020945.1 | 100 | 0 |
| Thermotogota | Athalassotoga saccharophila NAS-01 | GCA_009936215.1 | 96.61 | 0.62 |
| Thermotogota | Fervidobacterium pennivorans DSM 9078 | GCA_000235405.3 | 98.25 | 3.51 |
| Thermotogota | Thermosipho melanesiensis BI429 | GCA_000016905.1 | 98.28 | 0 |
| Thermotogota | Pseudothermotoga hypogea DSM 11164 | GCA_000816145.1 | 100 | 0 |
| Thermotogota | Thermotoga neapolitana DSM 4359 | GCA_000018945.1 | 100 | 0 |
| Thermotogota | Mesotoga prima MesG1.Ag.4.2 | GCA_000147715.3 | 98.12 | 0.47 |

| | | | | |
|---|---|---|---|---|
| Thermotogota | Kosmotoga pacifica SLHLJ1 | GCA_001027025.1 | 98.28 | 1.72 |
| Thermotogota | Marinitoga piezophila KA3 | GCA_000255135.1 | 98.28 | 1.72 |
| Thermotogota | Defluviitoga tunisiensis L3 | GCA_000953715.1 | 100 | 0 |
| Thermotogota | Petrotoga mobilis SJ95 | GCA_000018605.1 | 96.55 | 0 |

**Supplementary Data 1**

**Genome-scale metabolic model of *Zhurongbacter thermophilus* 3DAC (GEM-i3DAC).**

Eight sheets are included in Supplementary Data 1

| Sheet name | Description |
|---|---|
| 0-annotation_transcriptome_GPR | 3DAC genome annotations, transcriptomic data with or without sulfur in culture medium and the gene-reaction associations in 3DAC model. |
| 1-reactions | all reactions in 3DAC model |
| 2-compounds | all compounds in 3DAC model |
| 3-BOF | Biomass objective function formulation for biomass in 3DAC model |
| 4-exchange_constraints | Exchange constraints based on the culture medium in 3DAC model |
| 5-carbon_utilization | Simulations of carbon_utilization in 3DAC model, supporting the Fig. 2b |
| 6-electron_acceptor | Simulations on varied electron acceptors in 3DAC model, supporting the Fig. 2d |
| 7-gene_deletion | Simulations of the WT strain, single, double and triple deletions of MBS, Nfn2 and SH1, supporting the Fig. 2e |

**Supplementary Data 2**

**Genome-scale metabolic model of the last common ancestor of the superphylum Zhurongbacteria.**

Six sheets are included in Supplementary Data 2

| Sheet name | Description |
|---|---|
| 0-KO-list | Predicted KO list of the last common ancestor of superphylum Zhurongbacteria |
| 1-reactions | all reactions in the ancestor model |
| 2-compounds | all compounds in the ancestor model |
| 3-AA-biosynthesis | Statistics of the existing and missing reactions in biosynthesis pathways of all 20 amino acids based on KO of the ancestor |
| 4-exchange_constraints | Initial exchange constraints of the ancestor model |
| 5-simulations | Simulations of the utilization of different sole carbon sources in the ancestor model |

**Supplementary Data 3**

**Alignments and phylogenetic trees resulting from this study.**
